# Inhibition of Acyl-CoA Synthetase Long Chain Isozymes Decreases Multiple Myeloma Cell Proliferation and Causes Mitochondrial Dysfunction

**DOI:** 10.1101/2024.03.13.583708

**Authors:** Connor S. Murphy, Victoria E. DeMambro, Samaa Fadel, Heather Fairfield, Carlos A. Garter, Princess Rodriguez, Ya-Wei Qiang, Calvin P. H. Vary, Michaela R. Reagan

## Abstract

Multiple myeloma (MM) is an incurable cancer of plasma cells with a 5-year survival rate of 59%. Dysregulation of fatty acid (FA) metabolism is associated with MM development and progression; however, the underlying mechanisms remain unclear. Acyl-CoA synthetase long-chain family members (ACSLs) convert free long-chain fatty acids into fatty acyl-CoA esters and play key roles in catabolic and anabolic fatty acid metabolism. The Cancer Dependency Map data suggested that ACSL3 and ACSL4 were among the top 25% Hallmark Fatty Acid Metabolism genes that support MM fitness. Here, we show that inhibition of ACSLs in human myeloma cell lines using the pharmacological inhibitor Triascin C (TriC) causes apoptosis and decreases proliferation in a dose– and time-dependent manner. RNA-seq of MM.1S cells treated with TriC for 24 h showed a significant enrichment in apoptosis, ferroptosis, and ER stress. Proteomics of MM.1S cells treated with TriC for 48 h revealed that mitochondrial dysfunction and oxidative phosphorylation were significantly enriched pathways of interest, consistent with our observations of decreased mitochondrial membrane potential and increased mitochondrial superoxide levels. Interestingly, MM.1S cells treated with TriC for 24 h also showed decreased mitochondrial ATP production rates and overall lower cellular respiration.

**Implications:** Overall, our data support the hypothesis that suppression of ACSL in human MM cells inhibit their growth and viability, indicating that ACSL proteins may be promising therapeutic targets in treating myeloma progression.

## INTRODUCTION

Multiple myeloma (MM) is an incurable cancer of plasma cells with a 59% 5-year survival rate (1). A hallmark of cancer cells is their ability to adapt to their increased energy demands because of dysregulated cellular metabolism (2). Indeed, alterations in MM cell glucose and glutamine metabolism have been shown to be tumor-supportive, and thus underscore targets in metabolic pathways as promising preclinical therapeutic targets (3). However, targeting myeloma cell fatty acid (FA) metabolism as a novel anti-myeloma avenue has only been suggested by a limited number of preclinical studies (4–8).

Recent work on other cancers shows promise for targeting members of the acyl-coenzyme A long chain synthetase (ACSL) family of proteins, which activates long-chain fatty acids (saturated and unsaturated FAs with chain lengths of 8–22 carbons) into fatty acyl-CoA esters, which can then be used for catabolic or anabolic metabolism (9). In the catabolic pathway, FAs undergo fatty acid oxidation (FAO) in the mitochondria to generate ATP, whereas anabolically, FAs provide the substrates needed to synthesize triacylglycerols (TAGs, used to store energy), phospholipids, or other cell and organelle membrane lipids. System-level analyses of ACSLs across cancer types have revealed that ACSL expression and their role as oncogenes or tumor suppressors are heterogeneous and cancer-type dependent (10). While ACSL1 has been shown to support the proliferation of both colorectal (CRC) and breast cancer (BC) cell lines, evidence supports ACSL1 as a tumor suppressor in non-squamous cell lung carcinoma (NSCLC) cells (10,11). Additionally, ACSL1 and ACSL4 support invasion of CRC, prostate cancer, and quadruple-negative BC cells (12–15). In estrogen receptor-positive BC, ACSL4 has been shown to modulate drug efflux pumps to support chemotherapy resistance (13). ACSLs also regulate metabolism in a cancer type-specific manner. In CRC, overexpression of *ACSL1* and *ACSL4* resulted in enhanced glycolysis (11), while ACSL3 regulates FAO in BC and NSCLC in opposite directions, while supporting proliferation in both cancers (16,17). Therefore, it is critical to study the role of ACSLs in a cancer specific manner.

ACSLs play important roles in the development of lymphoid hematological malignancies. In a retrospective study of leukemia patients, *ACSL6* expression was positively correlated with overall survival, suggesting that ACSL6 may be a tumor suppressor in leukemia (10). In that analysis, of all ACSLs, only *ACSL4* was shown to be overexpressed in myeloma cells relative to control tissues (10). Zhang *et al.* also reported that *ACSL4* was overexpressed in primary MM cells and supported MM cell proliferation, possibly involving the c-Myc/sterol regulatory element binding protein (SREBP) axis (8). These investigators also showed that *ACSL4* expression in MM cells was positively correlated with their sensitivity to ferroptosis, an iron-dependent form of cell death (8). Despite these studies, the contribution of the ACSL family to multiple myeloma cell fitness remains largely unaddressed.

Triascin C (TriC) is an alkenyl-N-hydroxytriazene fungal metabolite and a competitive inhibitor of the FA-binding domain of ACSLs (18). TriC has been shown to effectively inhibit the growth of human breast cancer cells (19), while in acute myeloid leukemia (AML), TriC treatment inhibits cellular viability and synergizes with other anti-AML therapies (20). In endometrial cancer, TriC decreases survival, an effect that is enhanced by the addition of omega-3 FA docosahexaenoic acid (DHA) (21). TriC also induces apoptosis in glioma cells and synergizes with the apoptosis inducer etoposide, causing substantial cytotoxicity in glioma cells both *in vitro* and *in vivo* (19).

Overall, the data suggest that TriC is a safe, novel anticancer agent that has not been explored in myeloma cells. In the present study, we identified the ACSL family of enzymes as potentially supportive of MM cells using the Broad Institute’s Cancer Dependency Map database. We tested the hypothesis that ACSLs are MM supportive by quantifying the effects of TriC on human MM cell lines using *in vitro* functional and metabolic phenotyping assays. Using unbiased transcriptomic and proteomic analyses following TriC treatment of MM cells, we obtained new insights into the molecular phenotypic changes resulting from pan-ACSL inhibition in myeloma cells.

## MATERIALS AND METHODS

### Cancer Dependency Map Analysis

The gene fitness scores (Chronos scores) (22) for a modified list of the Hallmark Fatty Acid Metabolism genes (GSEA M5935 https://www.gseamsigdb.org, **Supp. Table 1**) in 21 human MM cell lines from the Cancer Dependency Map (DepMap) dataset (https://depmap.org/portal/download/) were reported. Human MM cell line gene and protein expression data were downloaded from the Cancer Dependency Map/Cancer Cell Line Encyclopedia (Expression 22Q2_Public) (23).

### Cell Lines and Culture Conditions

MM.1S (ATCC), RPMI-8226 (ATCC), and OPM-2 (DSMZ) cells, as well as luciferase– and gfp-expressing MM.1S (MM.1S^gfp+/luc+^) and MM.1R (MM.1R^gfp+/luc^) cell lines were obtained and cultured as previously described (7) in RPMI-1640 basal media supplemented with 10% (15% for U266B1 cells) fetal bovine serum (FBS, VWR) with 1% antibiotic-antimycotic (ThermoFisher Scientific, Cat # 15240112) at a cell density of 4.166×10^5^ cells/mL in tissue culture-treated T-75 flasks (VWR) (7). MM.1S^gfp+/luc+^ cells were used for experiments involving MM.1S cells unless otherwise stated. Human myeloma cell lines were authenticated by subjecting genomic DNA isolated with the QIAamp DNA Mini Kit (Qiagen), and short tandem repeat (STR) analysis was performed with the CLA IdentiFiler™ Plus PCR Amplification Kit (ThermoFisher Scientific) and sequenced on an ABI SeqStudio Genetic Analyzer (ThermoFisher Scientific) according to the manufacturer’s protocol through the Vermont Integrative Genomics Resource at the University of Vermont. STR profiles were compared between the experimental results and the reference using the Cellosaurus STR Similarity Search tool with the Tanabe algorithm, scoring only non-empty markers and excluding amelogenin (CLASTR v1.4.4, Swiss Institute of Bioinformatics).

### Triacsin C Treatment

Triacsin C (TriC) was purchased from Cayman Chemical (Ann Arbor, Michigan, USA). MM.1S^gfp+/luc+^, MM.1R^gfp+/luc+^, OPM-2, RPMI-8226, and U266B1 cells were seeded into either tissue culture-treated white bottom 96 well plate (4.33×10^4^ cells/well), tissue culture treated 24 well plates (1×10^5^ cells/well), tissue culture-treated 6-well dishes (4.81×10^5^ cells/well), or tissue culture-treated T-25 flasks (2×10^6^ cells/flask; Avantor/VWR, Cat. No. 10861-568) under the growth conditions described above. MM cells were treated with TriC or dimethyl sulfoxide (DMSO, vehicle). Samples were collected at 24-hour intervals and subjected to functional analyses below.

### Myeloma Cell Quantification, Viability and Apoptosis

For quantification and viability testing, MM cells were collected and resuspended in RPMI-1640 +10% FBS + 1% Anti/Anti, and diluted 1:2 in 0.4% Trypan Blue. Viable and non-viable cells were counted using a hemocytometer. To characterize apoptosis, MM cells were collected, washed 3 times with Cell Staining Buffer (BioLegend, Cat. No. 420201) and stained with APC-Annexin V (1:20, BioLegend Cat. no. 640920), DAPI (0.004 μg/ mL, ThermoFisher, Cat. No. D1306) in Annexin V Binding Buffer (BioLegend, Cat. no. 422201) for 15 min at room temperature). For all flow cytometric TriC analyses, a minimum of 1×10^4^ events were collected per sample on a MACSQuant Analyzer (Miltenyi Biotec) and analyzed using FlowJo v.10 (Becton, Dickinson & Company, Ashland, OR).

### Intracellular Characterization of BAX Protein

MM.1S cells were washed three times with Cell Staining Buffer (BioLegend) and then fixed in 1x Fixation Buffer (4% paraformaldehyde, BioLegend). Cells were washed 3 times with Cell Staining Buffer and stained with either Alexa Fluor (AF) 488 mouse anti-human BAX antibody (0.5 μg/mL, Biolegend, Cat. No. 633603) or AF488 Mouse IgG1 κ isotype control (5 μg/mL, Biolegend, Cat. No. 400129) in 1x Intracellular Staining Permeabilization Wash Buffer (Perm/Wash, BioLegend, Cat. No. 421002) for 15 min at room temperature. Cells were washed 2 times in 1x Perm/Wash buffer and resuspended in Cell Staining Buffer (Biolegend) prior to flow cytometry analysis. A total of 2×10^4^ events were collected using a MACSQuant (Miltenyi Biotec) and analyzed using FlowJo v10.6.1. Data are presented as the mean fluorescence intensity (MFI) of the FITC-H channel within the FSC-A and SSC-A gates.

### Myeloma Cell Cycle and Ki-67 Staining

MM cells were washed three times with Cell Staining Buffer (BioLegend) and fixed in 1x Fixation Buffer (4% paraformaldehyde, BioLegend). Cells were washed three times with Cell Staining Buffer and stained with Alexa Fluor 647 anti-human Ki-67 antibody (1:100) and DAPI (0.5 µg/ml) respectively in 1x Intracellular Staining Permeabilization Wash Buffer (BioLegend). The cells were resuspended in cell staining buffer (BioLegend, Cat. No. 420201) prior to flow cytometry using a MACSQuant Analyzer (Miltenyi Biotec).

### Flow Cytometric Characterization of Mitochondrial Number/Mass, Mitochondrial Membrane Potential, and Mitochondrial Superoxide Levels

MM cells were washed three times with Cell Staining Buffer (BioLegend) and resuspended in their respective cell culture media with 100 nM MitoTracker Green (Invitrogen, Cat. No. M7514), and incubated for 30 min at 37 °C. Cells were washed three times with cell staining buffer and resuspended in Cell Staining Buffer (BioLegend) prior to flow cytometry using a MACSQuant Analyzer (Miltenyi Biotec). To characterize mitochondrial membrane potential, MM cells were washed three times with Cell Staining Buffer (BioLegend) and resuspended in tetramethylrhodamine ethyl ester (TMRE) buffer (Cayman Chemicals) containing 100 nM TMRE (Cayman Chemicals, Cat. no. 701310) and incubated for 30 min at 37 °C. Cells were pelleted and resuspended in TMRE buffer and subjected to flow cytometry on a MACSQuant Analyzer (Miltenyi Biotec). For mitochondrial superoxide measurements, ATCC MM.1S cells were washed three times with cell staining buffer (BioLegend) and stained with 5 μM MitoSOX^TM^ Red (Invitrogen, Cat. No. M36008) in Hank’s balanced salt solution with calcium and magnesium (HBSS/Ca^2+^/Mg^2+^, Giboco, 14025-092) for 10 min at 37 C. Cells were washed three times with warm HBSS/Ca^2+^/Mg^2+^ and resuspended in HBSS/Ca^2+^/Mg^2+^ before analysis by flow cytometry.

### Cellular Metabolic Analysis

5×10^6^ MM.1S^gfp+/luc+^ cells were treated with DMSO or 1.00 μM TriC for 24 h in T-25 flasks (Avantor/VWR; Cat. no. 10861-568). Cells were then harvested, centrifuged, and resuspended in XF DEM media (pH 7.4; Agilent, Cat # 103575-100) containing 1mM sodium pyruvate, 10mM glucose and 2mM glutamine prior to plating on Seahorse XF 96 PDL-coated plates (Agilent, Cat # 103730-100) at a density of 75,000 cells/well per the manufacturer’s instructions. Mitochondrial function was determined using a Mitochondrial Stress Test on the Seahorse XFe96 analyzer (Agilent Technologies), as previously described (7). Cells were also analyzed for total, mitochondrial, and glycolytic ATP production rates using a Seahorse XF ATP Production Rate Assay according to the manufacturer’s instructions. The data presented here are representative of at least three independent experiments with ≥24 wells per treatment.

### Acyl-CoA Synthetase Long-chain Activity Assay

This protocol was adapted from Nchoutmboube *et al.* (2013) (24). MM.1S cells were plated in 6 well dishes (4.81×10^5^ cells/well) in RPMI+0.5% Fatty Acid (FA) Free BSA + 1% Anti/Anti and incubated with 0.5 μM BODIPY FL C16 at 37 C for 2 hours. The cells were incubated with TriC or DMSO for 2 h, collected and washed 3x with 0.2% FA-free BSA/PBS (ThermoFischer, Cat. No. AAJ6494422) to remove excess label. Cells were resuspended in 8.5% Sucrose + 0.5 μM EDTA + 10 mM Tris Buffer (pH 8.0) + 0.1% Triton X-100 and incubated at room temperature for 25 min. Lysates were centrifuged for 10 min at 14,000 × g, and the supernatant was transferred to fresh tubes. Heptane was added to the supernatant (1 volume of supernatant: 6 volumes of n-heptane), shaken at 1,300 rpm for 10 min, and centrifuged for 5 min at 12,000 × g. n-Heptane was removed using a pipette, and the aqueous layer was subsequently extracted with n-heptane three more times. The aqueous layer was read on a black 96-well plate (Corning Cat No. 3603, 475 nm /500-525 nm, excitation/emission). Heptane (Cat. no. 34873) was purchased from Millipore Sigma (Burlington, MA, USA).

### Total mRNA extraction and quantitative real-time polymerase chain reaction (qRT-PCR)

Total RNA was harvested in QIAZOL and prepared using the Qiagen miRNEASY Kit with DNase On-column digestion (Qiagen, Hilden, Germany), according to the manufacturer’s protocol. mRNA was quantified and tested for quality and contamination using a Nanodrop 2000 (Thermo Fisher Scientific) and subjected to quality control minimum standards of 260/230>2.0 and 260/280>1.8 before downstream applications. For qPCR, cDNA was synthesized using MultiScribe reverse transcriptase (High-Capacity cDNA, Applied Biosciences, ThermoFisher Scientific) according to the manufacturer’s instructions using 500 ng of total RNA. Relative transcript expression was determined using SYBR Master Mix (Bio-Rad, Hercules, CA) and thermocycling reactions on a CFX-96 or Opus system (Bio-Rad) using 500 ng of cDNA. Target transcripts (**Supplementary Tables 1, 2**) were normalized to *TATA-box binding protein* (*TBP)* using the 2^-ΔΔct^ method. Data were analyzed using Bio-Rad CFX Manager 3.1.

### RNA Sequencing Sample Preparation and Analysis

A total of 5 × 10^6^ MM.1S^gfp+/luc+^ cells were seeded in T-25 flasks (Avantor/VWR, Cat. No. 10861-568) and treated with the vehicle (DMSO) or 1.00 μM triacsin C for 24 h. Replicates were defined as MM.1S^gfp/luc+^ cells of the same passage grown in parallel. After 24 h, RNA was isolated using a Qiagen RNeasy Plus Mini Kit (Qiagen, Hilden, Germany) Cat. no. 74136), according to the manufacturer’s protocol. Samples were evaluated on Bioanalyzer BA210O RNA pico chips and quantified using the Qubit HS DNA reagent. Sequence libraries were prepared with the Takara Pico V2 library prep using 6 ng of total RNA and sequenced with an Illumina HiSeq 1500/2500. RNA sequence data were analyzed using the nf-core/rnaseq pipeline v3.9 using the Nextflow workflow manager v22.10.2. Raw reads were subjected to quality checking and reporting (FastQC v0.11.9/ MultiQC v1.13), and low-quality sequence data (Phred score <20) were removed using Trim Galore v 0.6.7. The reads were aligned to the *Homo sapiens* hg38 reference genome using STAR v2.7.10a and SAMtools v1.15.1. Read counts were quantified using SALMON v1.5.2, and DESeq2 v1.28.0 was used to identify differentially expressed genes using a cut-off value of (log2(FC) > |1|, q-value < 0.05) using the Wald test and adjusted for multiple testing using the Benjamini and Hochberg method. Gene ontology enrichment analysis was performed using the Enrichr package and STRINGv11, with a high confidence score cutoff of 0.70.

### Sample Preparation for Mass Spectrometry Proteomics of TriC treated MM.1S cells

MM.1S^gfp+/luc+^cells were treated with vehicle (DMSO) or 1.00 or 2.00 μM triacsin C for 48 h. Cells were collected and washed 3x with cold PBS, and flash-frozen. Cells were solubilized in ice-cold RIPA buffer, and DNA was sheared using a probe-tip sonicator (Branson Ultrasonifier 250, Branson Ultrasonic Corporation, Danbury, CT, USA, 3 × 10 s). Each sample was then centrifuged (14,000 × g) at 4°C and the supernatant was collected. One hundred micrograms of protein was taken from each sample and reduced using 5 mM TCEP (tris(2-carboxyethyl)phosphine hydrochloride; Strem Chemicals, Newburyport, MA). The reaction was allowed to proceed for 20 min at 56°C and alkylated for 30 min in the dark with 10 mM iodoacetamide at room temperature (G-Biosciences, St. Louis, MO, USA). Proteins were precipitated at –20°C with ethanol, and pellets were washed twice with ice-cold ethanol followed by overnight incubation at 37°C in 100 mM ABC containing 1 mM CaCl_2_ and trypsin (Sequencing grade, modified, Promega Co, Madison, WI, USA). Digested proteins were evaporated and each sample was freed from salts and buffers by solid-phase extraction on C18 resin using cartridges prepared in-house. Briefly, for each sample, a C18 StageTip was prepared according to the previously reported procedure (25). Octadecyl-derivatized silica (4 mg; SiliaSphere PC, C18 monomeric, 25 µm particles, 90 Å pore size, SiliCycle Inc., Québec City, Canada) suspended in LC-MS-grade isopropanol (Honeywell, Morris Plains, NJ, USA) was added to each tip. Each cartridge was then equilibrated, and the samples were purified according to the StageTip protocol referenced above. Purified peptides were eluted directly into autosampler vials for use in LC-MS instrumentation using 100 µL elution buffer, and the solvent was removed by vacuum centrifugation. Each sample was then resuspended in a volume of sample LC/MS loading solvent [5% formic acid (Optima grade, ThermoFisher Scientific) and 4% acetonitrile (both water and acetonitrile were LC-MS-grade, Honeywell)] to yield an approximate concentration of 1 µg/µL peptides.

### Mass Spectrometry Proteomics of TriC-treated MM.1S Cells

Chromatography, mass spectrometry, and data analysis were performed as previously described (26). Key differences between the protocols are highlighted here. Sample separation was performed on an Eksigent NanoLC 425 nano-UPLC System (Sciex, Framingham, MA, USA) in direct-injection mode with a 5 µL sample loop made in-house. The analysis was performed in positive ion mode on a TripleTOF 5600 quadrupole time-of-flight mass spectrometer (Sciex).

### Data Availability Statement

Raw and normalized RNA-Seq data are available from the Gene Expression Omnibus database (GSE252929). Mass spectrometry data are available in the PRIDE database, PXD049304.

### Statistical Analysis

All graphs were created using GraphPad Prism (v9 or above); statistical significance was determined using One-way or Two-way ANOVA with Tukey’s, Šídák’s, or Dunnett’s multiple comparisons tests, Student’s t-test, or Welch’s test unless otherwise stated. Data represent the mean ± standard deviation, unless otherwise noted. Significance is indicated as: *, p<0.05; **, p<0.01; ***, p<0.001; **** p<0.0001.

## RESULTS

### Pharmacological Inhibition of the Long-Chain Acyl-CoA Synthetase (ACSL) Family Decreases Multiple Myeloma Cell Proliferation and Survival

To test the hypothesis that fatty acid metabolism supports multiple myeloma proliferation and survival, we investigated the Cancer Dependency Map (DepMap) (22), a collection of fitness scores of a set of human multiple myeloma cell lines containing CRISPR/Cas9 knockouts from the Hallmark Fatty Acid Metabolism gene set (Molecular Signature Database: M5935) (27). We observed that four of the five human **a**cyl-**C**oA **s**ynthetase **l**ong chain family members (ACSL) had negative Chronos scores, indicating that *ACSL3* and *ACSL4* were among the top 25% most essential fatty acid metabolism genes in MM cells (**Figure 1A, Supp Table 3**). These data suggest that *ACSL1, ACSL3, ACSL4* and *ACSL5* support human multiple myeloma cell growth and survival. To better understand the landscape of ACSL gene and protein expression in MM cell lines, we consulted the DepMap. The average gene expression of *ACSL1*, *3*, *4* and *5* ranged from 3.78-5.19 TPM+1, with *ACSL3* having the highest average expression in the family and *ACSL6* having the lowest (0.05 TPM+1) (**Supp.** Figure 1A). The DepMap ACSL protein expression data from six human myeloma cell lines showed varied expression of each ACSL family member between each MM cell line (**Supp Figure 1B**).

**Figure 1:**
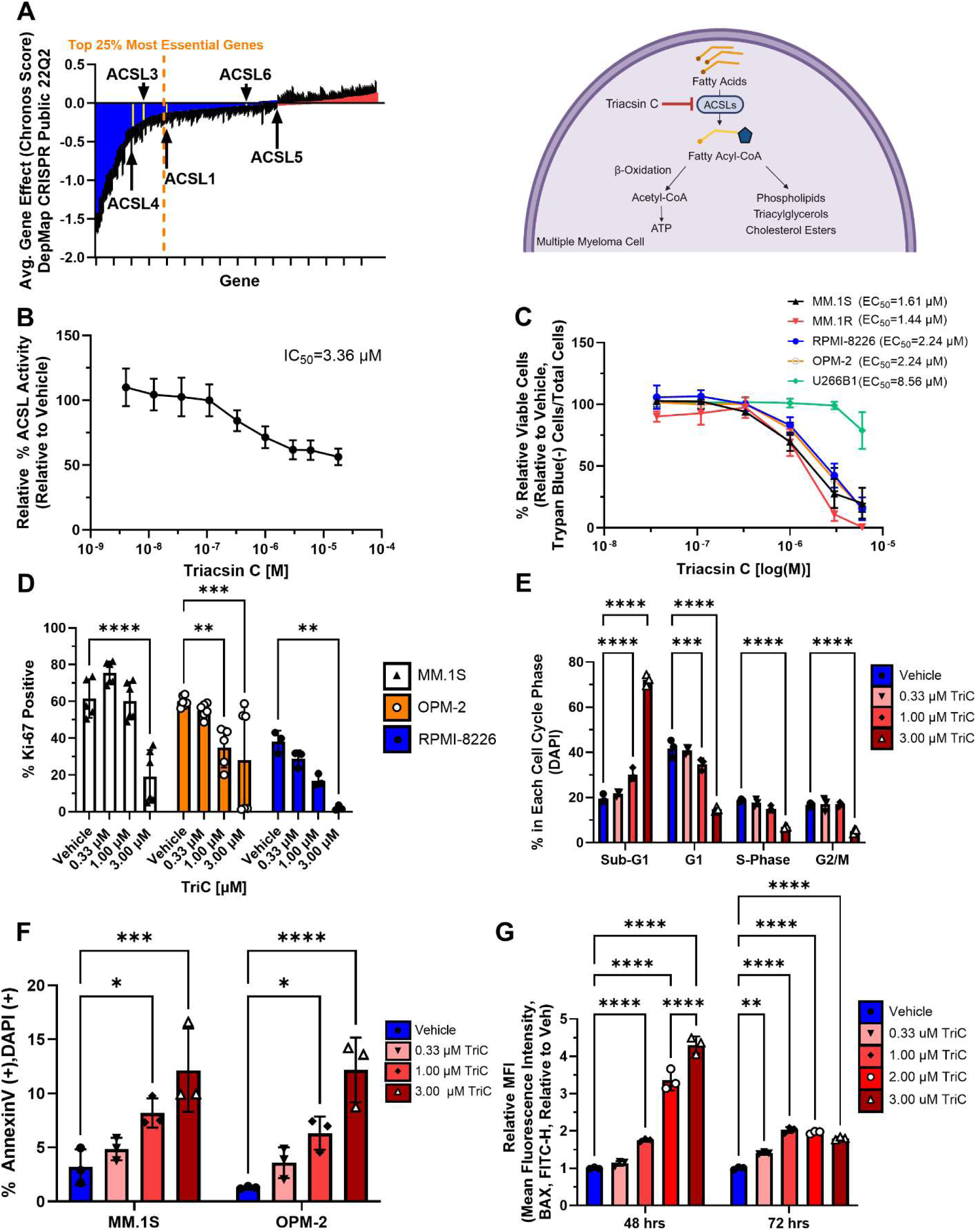
Targeting the ACSLs using Triacsin C Inhibits Myeloma Cell Proliferation and Survival. **A**, Hallmark Fatty Acid Metabolism genes from Gene Set Enrichment Analysis (M5935) displayed as the average gene fitness (Chronos Score) data from the Cancer Dependency Map of of 21 human myeloma cell lines. Blue bars represent targets of interest with average Chronos scores < 0.0 (myeloma cell growth fitness defect upon CRISPR knockout), while red bars are Chronos scores >0.0. Schematic was made with Biorender.com and adapted from Tang *et al.* 2018, *Oncology Letters* (50); ACSL family members are highlighted in yellow. **B,** ACSL activity relative to vehicle treated cells of ATTC MM.1S cells treated with various doses of triacsin C (TriC) for 2 hours **C.** MM.1S, MM.1R, RPMI-8226, OPM2 and U266B1 cells were incubated with various doses of TriC for 48 hrs and stained with Trypan Blue to quantify viable cells/mL. EC_50_ values were calculated using Non-linear regression (four parameter, variable slope) in GraphPad Prism v9.4.1. n=3 **D,** Proliferation of human MM cell lines MM.1S, OPM-2 and RPMI-8226 cells treated with various doses of TriC for 48 hrs and stained with AF647 anti-human Ki67 (% positive); n=3-6. **E,** Cell cycle distribution of MM.1S cells treated with various doses of TriC for 48 hrs and stained with DAPI; n=3. **F,** Apoptosis assay using Annexin V/DAPI staining of MM.1S, U266B1 and OPM2 cells treated with various doses of TriC for 48 hrs; n=3. **G,** Intracellular BAX (AF488 mouse anti-human BAX antibody) levels in MM.1S cells treated with various doses of TriC for 48 hrs; data displayed as FITC-H MFI; n=3. Significance determined via two-way ANOVA with Statistics: Tukey’s multiple comparison test **(D-F)** or Šídák’s multiple comparisons test **(G)** All data are mean ± StDev, *p<0.05, **p<0.01, ***p<0.001 ****p<0.0001

Triacsin C, a small molecule inhibitor of ACSL1,3,4 and 5 (28), was used to test the hypothesis that the ACSL family supports MM cell fitness. To determine the IC_50_ of TriC and a rational dose range for subsequent experiments, we measured the total cellular ACSL activity (24). Briefly, MM.1S cells were incubated with a fluorescent fatty acid substrate BODIPY FL C_16_ for 2 h, challenged with various doses of triacsin C for 2 h, and subjected to n-heptane extraction to separate the product (BODIPY FL C_16_) from the substrate (BODIPY FL C_16_-CoA). The IC_50_ of triacsin C was 3.66 μM (**Figure 1B**). To test the hypothesis that ACSL family inhibition by triacsin C would decrease MM cell proliferation and survival, human MM cell lines (MM.1S, MM.1R, RPMI-8226, OPM-2, and U266B1) were treated with TriC (0.0366-6.00 μM) at 24-hour intervals for up to 72 h and subjected to trypan blue staining to assess cell viability. Significant differences in MM cell viability were first observed after 48 hours of TriC treatment with most human MM cells responding with a dose-dependent decrease in viability (average EC_50_:1.88 μM), except for U266B1 cells (EC_50_:8.56 μM) (**Figure 1C, Supp Figure 1C**).

To test the contributions of cytotoxic and cytostatic effects of TriC, we measured MM cell proliferation via Ki-67 staining, cell cycle (DAPI), and apoptosis by Annexin V/DAPI staining, and by measuring BAX protein levels for 72 hours at 24-hour intervals. At 48 h, MM.1S, OPM-2 and RPMI-8226 cells showed a significant decrease in Ki-67 (+) cells at 3 μM TriC (**Figure 1D**). Interestingly, treatment with 1 and 3 μM TriC for 48 h significantly decreased the percentage of cells in G_1_ and G_2_/M phases and increased the population of sub-G_1_ MM.1S, OPM-2, and RPMI-8226 cells. As sub-G_1_ cells are associated with DNA degradation, these data suggest that TriC induces DNA fragmentation but not cell cycle arrest (**Figure 1E, Supp.** Figure 1F,G). Consistent with these data, apoptotic cells [Annexin V(+), DAPI(+)] increased (7-12% total apoptosis) in a dose-dependent manner at both 48 and 72 h post-TriC treatment in MM.1S and OPM-2 cells (**Figure 1F**). Moreover, we observed a dose-dependent increase in the key apoptotic protein BAX in MM.1S cells treated with TriC for 48 and 72 h (**Figure 1G**). Taken together, these data demonstrate that TriC treatment decreases the viability and proliferation of human MM cell lines after 48 h of treatment and is characterized by an increase in the percentage of sub-G_1_ cells, increases in Annexin V/DAPI-positive cells, and the apoptosis-related protein BAX.

### Triacsin C Induces Transcriptional Changes in MM.1S Cells Associated with Cell Death and the Integrated Stress Response

In order to understand the mechanism underlying TriC toxicity in MM cells, MM.1S cells were treated *in vitro* with vehicle or 1 μM TriC for 24 h and subjected to RNA-seq analysis. Transcripts for each sample were aligned to the *Homo sapiens* GRCh38 genome, which included 12,772 protein-coding genes, all of which passed the read mapping and quality parameters (**Supp.** Figure 2C). Principal component analysis and comparison of Euclidean distances between samples showed that the transcriptomes of vehicle– and TriC-treated cells were distinct (**Figure 2A and Supp.** Figure 4A). Between TriC and vehicle-treated MM.1S cells, DESeq2 analysis revealed 208 differentially expressed genes (DEGs; log2(FC) > |1|, padj < 0.05) (29) which included 167 upregulated and 41 downregulated genes (**Figure 2B**). Enrichr was used to identify significantly enriched pathways within the 208 DEGs. Significant pathways of interest in both the Reactome (30) and KEGG (31) pathways of upregulated DEGs included: Cellular Response to Stress, ATF4 Activation in Response to Endoplasmic Reticulum Stress, Ferroptosis, and Apoptosis (**Figure 2C,D, Supp. Table 4, 5**). Notable Reactome and KEGG pathways enriched among the significantly downregulated genes in TriC-treated cells were Aryl Hydrocarbon Receptor Signaling and POU5F1 (OCT4), SOX2, and NANOG activated genes related to proliferation (**Supp.** Figure 4E,F**, Supp. Table 6, 7**).

**Figure 2:**
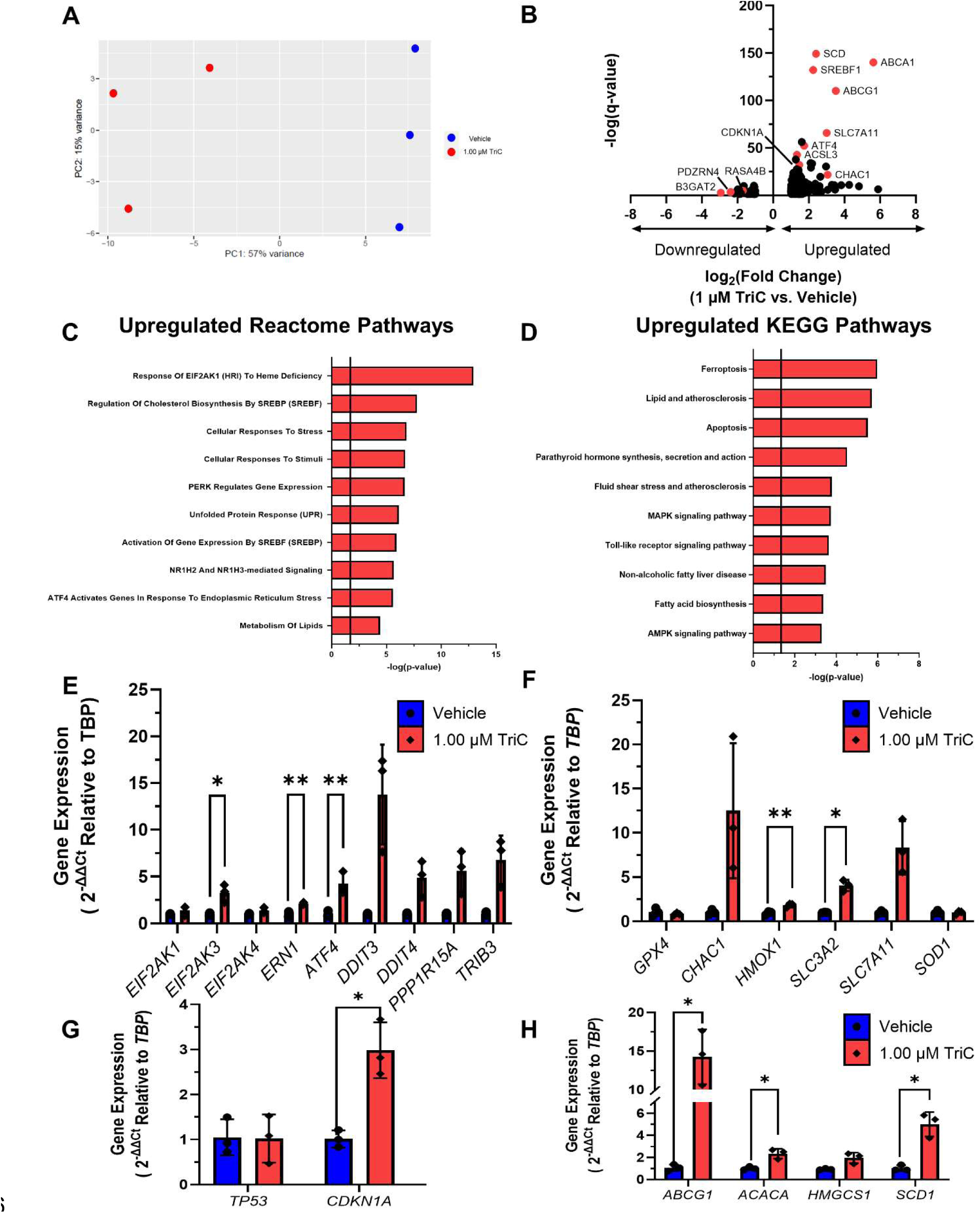
Triacsin C Treatment of MM.1S Cells Induces Transcriptional Changes Associated with Cell Death and the Integrated Stress Response. **A**, Transcriptional profiles of MM.1S cells treated with vehicle (DMSO) or 1 μM TriC for 24 hours as assessed by PCA of RNA-Seq data. **B,** Differentially expressed transcripts derived from RNA-seq of MM.1S cells treated with vehicle (DMSO) or 1 μM TriC for 24 hours. **C-D,** Reactome and KEGG pathways (respectively) enriched in the significantly upregulated transcripts in 1 μM TriC treated MM.1S as determined via Enrichr. **E-H,** Expression of genes related to *ATF4* signaling (**E**), ferroptosis (**F**) *TP53* signaling (**G**) and fatty acid metabolism (**H**) relative to *TBP* in MM.1S cells treated with vehicle or 1 μM TriC for 24hrs; n=3. **Statistics:** Significance was tested in panels **2E-H** with an unpaired Student’s t-test or Welch’s t-test. All data are mean ± StDev, *p<0.05, **p<0.01, ***p<0.001 ****p<0.0001

Gene overrepresentation analysis suggests that TriC activates EIF2AK3-EIF2S1-ATF4 signaling and cell death pathways and negatively regulates proliferation. Indeed, we observed significantly increased gene expression of *EIF2AK3* (3.2 fold) and *ATF4 (*4.2 fold) and a trend of increased expression in a number of its downstream targets, such as *PPP1R15A* (5.6 fold)*, TRIB3* (6.7 fold) and the pro-apoptotic DDIT3 (13.7 fold) in MM.1S cells treated with 1 μM TriC relative to vehicle-treated cells. Furthermore, there was a significant increase in a key gene in another arm of the ER stress pathway, *ERN1* after 24 h of treatment with 1 μM TriC in MM.1S cells. There were no significant changes in gene expression of other kinases that activate EIF2S1, *EIF2AK1* or *EIF2AK4*. (**Figure 2E**). In addition to EIF2AK3-EIF2S1-ATF4 activation, a gene associated with mitigating ferroptosis, *SLC3A2,* (4.0 fold) was significantly increased, with a trending increase in gene expression of the related *SLC7A11*. Additionally, the pro-ferroptotic genes, *HMOX1* (1.8 fold) had significantly increased expression and *CHAC1 (*13.9 fold) had an increasing trend in gene expression upon TriC treatment (**Figure 2F**). Moreover, the key tumor suppressor *CDKN1A* (2.9 fold) was significantly increased in TriC-treated MM.1S cells, with *TP53* levels remaining unchanged (**Figure 2G**). There was a significant (p<0.05) increase in genes involved in fatty acid metabolism, such as *ACACA* (2.3 fold)*, SCD1* (5.0 fold), and the cholesterol transporter *ABCG1* (14.2 fold) with a trending increase in the rate-limiting enzyme for cholesterol biosynthesis, *HMGCS1* (**Figure 2H**). Interestingly, there was a decrease in the gene expression of metastasis-associated, RAC1 (*PAK6,* 0.5 fold), and oncogenic RAS pathways (*RASA4B,* 0.2 fold) (**Supp.** Figure 4F). Thus, these data show that MM.1S cells treated with triacsin C for 24 h have a transcriptional profile associated with ATF4 activation, apoptosis, ferroptosis, and negative regulation of cell cycle progression.

### Triacsin C Treatment Induces Proteomic Changes Associated with Mitochondrial Dysfunction and Reactive Oxygen Species Detoxification

To identify global protein changes induced by TriC treatment, we treated MM.1S cells with 1 μM TriC, 2 μM TriC, or vehicle for 48 h and subjected them to sequential window acquisition of all theoretical fragment ion spectra (SWATH) mass spectrometry. The proteome of each treatment group was well-defined and functionally distinct from that of the other treatments, as assessed by PCA (**Figure 3A**). Of the approximately 1,580 total proteins detected in TriC-treated MM.1S cells, 167 and 614 differentially expressed proteins were detected in the 1 and 2 μM TriC-treated cells, respectively. The majority of differentially expressed proteins in both TriC treatments decreased (81.4% and 61.7%, 1 μM, and 2 μM, respectively) relative to vehicle-treated control cells (**Figure 3B,C**). The canonical pathway features in the Ingenuity Pathway Analysis (32) revealed six shared pathways among MM.1S cells treated for 48 h with both TriC doses, including: phagosome maturation, protein ubiquitination, FAT10 signaling, mitochondrial dysfunction, oxidative phosphorylation, and EIF2 signaling (**Figure 3D, E**). Interestingly, mitochondrial function and EIF2 signaling were predicted to be activated, while oxidative phosphorylation was likely inactivated (Z score ≥ |2|) based on the differential protein expression of both the 1μM and 2 μM TriC conditions compared to the vehicle (**Figure 3D, E**). Within the six shared dysregulated pathways, 39 differentially expressed proteins were common to both TriC doses tested in MM.1S (**Figure 3F, Supp. Table 8**), and these were significantly enriched for Biological Processes (identified using Enrichr) in two major categories: reactive oxygen species metabolism (SOD1, PRDX1, 2, 5, and 6) and mitochondrial electron transport (COX6B1, COX5A and COX7A2) (**Figure 3G**). Indeed, qRT-PCR gene expression of *COX5A*, a subunit of the genes of Complex IV, was significantly decreased in MM.1S cells treated with TriC for 48 h, with similar but non-significant trends for decreased expression of *COX6B1* and *ATP5ME* (**Figure 3H**).

**Figure 3:**
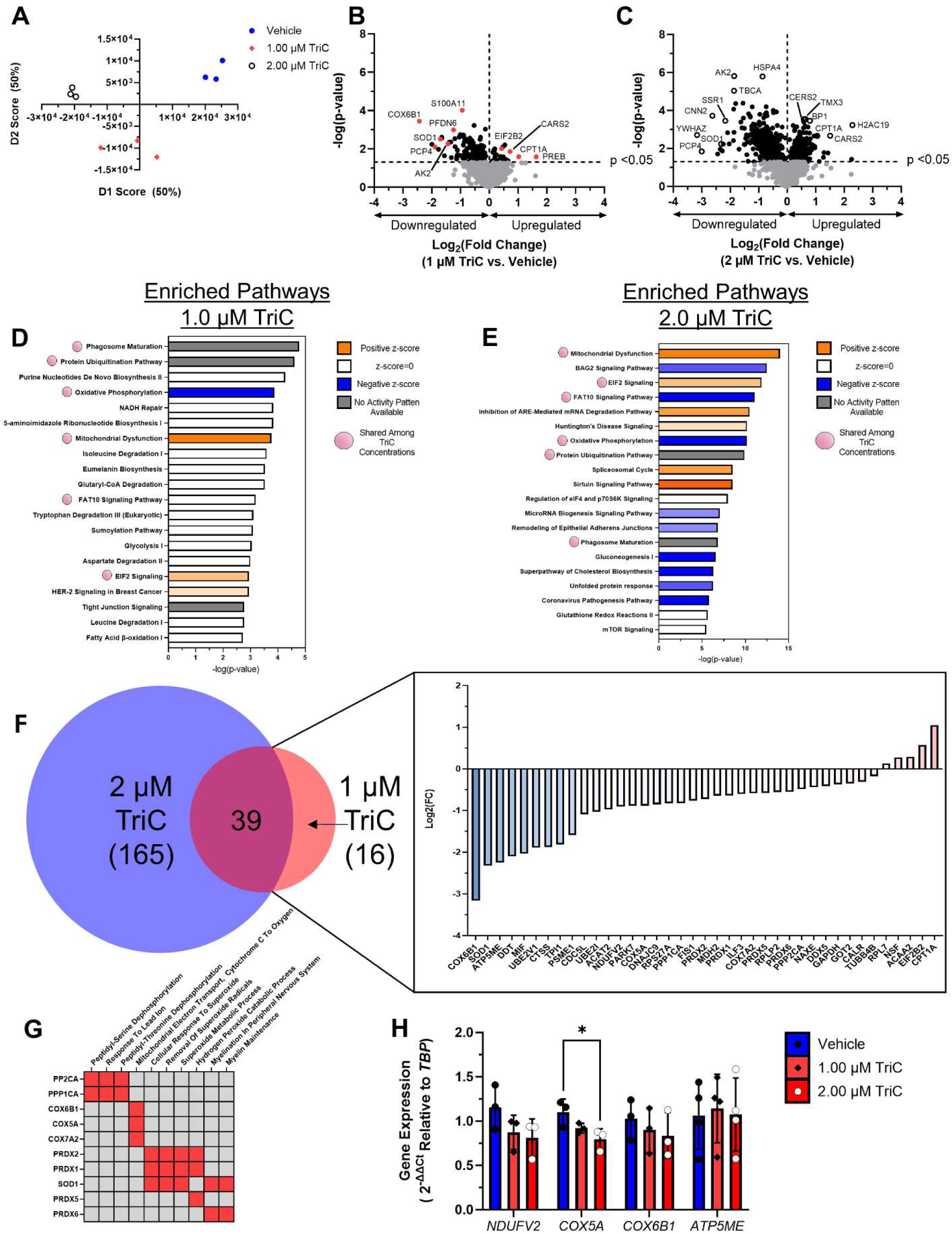
Triacsin C Treatment of MM.1S Cells Induces Proteomic Changes Associated with Mitochondrial Dysfunction and Reactive Oxygen Species. **A**, Proteomic profiles of MM.1S cells treated with Vehicle (DMSO, blue filled circles), 1 μM (red diamonds) or 2 μM (open black circles) TriC as assessed by principal component analysis. **B-C,** Aberrantly expressed proteins in MM.1S cells treated for 48hrs w/ either 1 or 2 μM TriC **D-E,** Top 20 enriched pathways in MM.1S cells treated with 1 and 2 μM TriC, respectively, identified using Ingenuity Pathway Analysis with their associated –log(p-value). **F,** DeepVenn depiction of the number of significantly changed proteins among the shared significantly changed pathways identified by IPA between 1 μM TriC (red, 16 unique proteins) and 2 μM TriC (blue, 165 unique proteins) with a total of 39 shared proteins (purple). The log_2_(fold change) is depicted for the 39 shared proteins among MM.1S^gfp+/luc+^ treated for 48 hrs with 1 or 2 μM TriC. **G,** Gene ontology (GO) enrichment for GO Biological Process of the 39 shared dysregulated proteins between MM.1S cells treated with 1 μM and 2 μM TriC treated for 48 hrs as assessed by Enrichr. A selection of proteins with common aberrant expression in the presence of both TriC doses are depicted here with red boxes indicating association between the protein and the GO term; gray boxes indicate no association. **H,** Expression of genes related to oxidative phosphorylation in MM.1S cells treated with vehicle or 1 µM TriC for 24hrs, as assessed by qRT-PCR; n=3. **Statistics:** Two-way ANOVA with Šídák’s multiple comparisons test (**H**) and Student’s t-test to identify differentially expressed proteins (**B-C**). All data are mean ± StDev, *p<0.05, **p<0.01, ***p<0.001 ****p<0.0001.

### Triacsin C Treatment Negatively Impacts Multiple Myeloma Cellular Metabolism and Mitochondrial Function

Given that TriC-treated MM.1S cells exhibited changes in mitochondria-related pathways (oxidative phosphorylation and mitochondrial dysfunction), we aimed to test the hypothesis that TriC impairs MM cellular metabolism and mitochondrial function. To assess the effect of TriC on MM cellular metabolism, we treated MM.1S cells with 1 μM TriC (MM.1S-TriC) for 24 h and then subjected the cells to mitochondrial stress measurements (**Figure 4A**). MM.1S-TriC cells had significantly reduced basal, maximal, and ATP-dependent mitochondrial respiration and proton leakage (**Figures 4A, B**). Interestingly, in parallel samples, we observed a significant 21.27% decrease in total ATP production rates attributable to a 57.3% decrease in mitochondrial ATP production rates, with no significant compensatory increase in glycolytic ATP production rates in MM.1S-TriC compared to MM.1S-Veh (**Figure 4C**). Taken together, these data demonstrate that TriC has profound effects on mitochondrial ATP production rates that reduce cellular respiration in MM.

**Figure 4.**
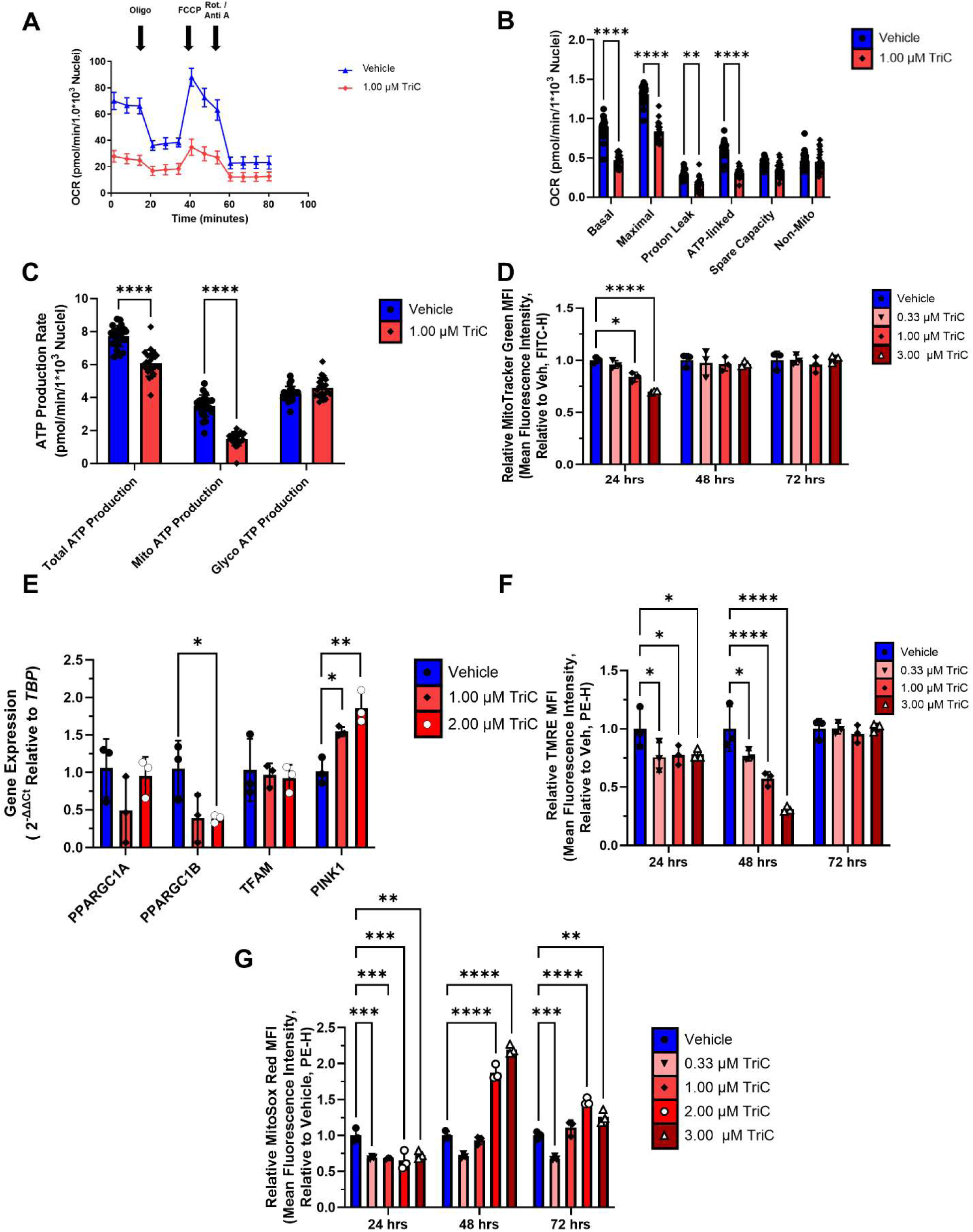
Triacsin C Induces Mitochondrial Dysfunction in MM Cells: **A-B**, Cellular respiration (oxygen consumption rate, OCR) in MM.1S cells treated with 1 μM triacin C (TriC) for 24 hours and subjected to a Mitochondrial Stress test. Values are normalized to the number of nuclei. Data are representative of 3 independent experiments. Oligo=oligomycin (Complex V inhibitor), FCCP (carbonyl cyanide p-trifluoro methoxyphenylhydrazone, proton gradient uncoupler), Rot/Anti A=Rotentone/antimycin A (Complex I and III inhibitors, respectively. **C,** Total, Glycolytic, and Mitochondrial ATP Production Rates in MM.1S cells treated with 1 μM TriC. **D,** Number/Mitochondrial Mass assessed with MitoTracker Green in MM.1S cells treated with TriC for 24, 48 and 72 hrs; n=3. **E,** Expression of genes related to mitochondrial biogenesis MM.1S cells treated with vehicle, 1.00 or 2.00 μM triacsin C for 48 hours; n=3. **Statistics:** Assessment of mitochondrial integrity in MM.1S cells were treated with TriC for 24, 48 and 72 hrs with TMRE staining; n=3. **F,** Mitochondrial superoxide quantification (MitoSox Deep Red) in MM.1S cells treated with TriC for 24, 48 and 72 hrs; n=3. **Statistics:** Two-way ANOVA with Tukey’s multiple comparison test used throughout except for **4E**, where a One-way ANOVA with Dunnett’s multiple comparisons test was used. Data are mean ± StDev *p<0.05, **p<0.01, ***p<0.001 ****p<0.0001

To test whether TriC alters mitochondrial biogenesis, total mitochondria were quantified by flow cytometry using MitoTracker Green staining **(Figure 4D).** TriC-treated MM.1S cells exhibited a dose-dependent decrease in the relative mean fluorescence intensity (MFI) of total mitochondria in MM.1S-TriC vs. MM.1S-Veh cells after 24 h, but not at 48 or 72 h of treatment. MM.1S cells treated with either 1 μM or 2 μM TriC after 48 h showed significant increases in gene expression of the key mitochondrial biogenesis gene, *PPARGC1B* relative to vehicle-treated cells, as assessed by qRT-PCR **(Figure 4E).** These data suggest that there is a decrease in total mitochondrial number after 24 h of treatment with triacsin C in MM.1S cells, but that this decrease was recovered over time. We next tested the mitochondrial membrane potential of MM.1S cells treated with TriC at 24-hour intervals for a total of 72 h using tetramethylrhodamine ethyl ester (TMRE). Consistent with the proteomic enrichment of mitochondrial dysfunction, we observed a dose-dependent decrease in the relative MFI of TMRE in MM.1S-TriCvs. MM.1S-Veh after 24 and 48 h of treatment. These data suggest that TriC induces mitochondrial dysfunction through the loss of mitochondrial membrane potential **(Figure 4F)**.

Mitochondrial dysfunction is often associated with an increase in oxidative stress. Coupled with our observation that the reactive oxygen species metabolism was an enriched biological process in the shared proteins among 1 and 2 μM TriC-treated MM.1S cells, we predicted that TriC-treated MM cells would exhibit increased ROS. Interestingly, MM.1S cells treated with TriC showed an initial decrease in mitochondrial superoxide by MitoSOX Deep Red staining MFI after 24 h of TriC treatment. However, treatment for 48 and 72 h resulted in a dose-dependent increase in the mitochondrial superoxide levels (**Figure 4G**). Taken together, TriC-treated MM cells showed compromised oxygen consumption, mitochondria-derived ATP production rates, decreased mitochondrial mass and mitochondrial membrane potential, and increased production of superoxide.

## DISCUSSION

Accumulating evidence suggests the potential of targeting the long-chain fatty acyl-CoA synthetase family in cancer (9). However, studies in multiple myeloma have been limited to ACSL4 (8). In this study, we report that within the Cancer Dependency Map’s genome-wide CRISPR/Cas9 screen, the majority of acyl-CoA synthetase long chain family members are supportive of human multiple myeloma cell fitness. Consistent with the hypothesis that ACSLs support MM cell proliferation and survival, we showed that pharmacological inhibition of the ACSL family with triacsin C in human MM cell lines decreased MM cell viability and proliferation and induced apoptosis starting after 48 h of treatment. These results are consistent with reports of TriC treatment decreasing viability in human breast cancer (33), Burkitt’s lymphoma (34), and endometrial (35) cell lines, and initiating apoptosis in human *TP53*-mutant lung, colon, and brain cancer cell lines (36). Interestingly, while most of the MM cell lines tested were *TP53* mutants, MM.1S (*TP53*^WT^) cells were sensitive to TriC treatment, suggesting that either TriC toxicity in MM cells is independent of *TP53* status, or that toxicity occurs via multiple mechanisms. We observed heterogeneity in the levels of Ki-67 positive cells treated with 3 μM TriC, which may be due to a change in the rate of decay of Ki-67 in cells undergoing cell death (37), and not all the ACSLs contributed equally to proliferation.

We initially found that human myeloma cells exhibited substantial basal expression of *ACSL1*, *ASCL3*, *ACSL4*, and *ACSL5*, albeit with minimal expression of *ASCL6*, suggesting that broad targeting by pharmaceutical methods could be impactful. Our calculated IC_50_ of TriC (3.36 μM) in human myeloma cells was consistent with other reports. In other systems, biochemical inhibition of ACSLs from rat liver homogenates found the concentrations of TriC required for 50% inhibition to be 8.7 μM (28). Similarly, in enzymatic activity studies using rat recombinant ACSL1 and ACSL4, the IC_50_ values for TriC were found to be 4–6 μM (38), and rat ACSL5 activity was found to be insensitive to TriC (38). The effectiveness of TriC may be species-specific, as TriC inhibits the growth of human BC cells and human intestinal cells, where *ACSL5* is highly expressed (19,39). TriC has also been shown to inhibit the viability of acute myeloid leukemia (AML) cell lines and primary cells (in part by inducing apoptosis), synergize with other anti-AML therapies, and show no toxicity to healthy donor CD34+ cord blood cells in the dose range tested (up to 16 µM, 48 h) (20). Future studies with TriC in MM designed to test EC_50_ in non-malignant plasma cells are needed to provide evidence that the toxicity of targeting the ACSL family is specific to MM cells. We found that the human MM cells tested were similarly sensitive to TriC, with EC_50_ values ranging from 1.44 µM (MM.1R) to 8.56 µM (U266B1) and far lower than reported doses toxic to non-cancerous cells (20). Importantly, cells treated with 1 µM TriC exhibited significantly higher levels of apoptosis and a shift from active cell cycle phases into sub-G1 compared to vehicle-treated cells, suggesting that inhibition of ACSLs in myeloma may be a plausible therapeutic target. Given the higher EC_50_ in TriC-treated U266B1 cells, it is possible that molecular subtypes of MM cells are intrinsically resistant to the effects of ACSL family inhibition. To address the possibility of TriC-dependent off-target effects and address concerns of compensation from ACSL isozymes after TriC treatment, future studies using genetic approaches to knockdown or knockout all ACSL isozymes and each individual ACSL isozyme in MM cells are warranted.

Distinct mechanisms of action have been implicated in the effects of TriC in other cells, including the reduction of key survival pathways (p38/MAPK (40), nuclear factor-κB (NF-kB) (40)), PPARy (41) levels, and increased Bax-induced caspase activation (19). In MM.1S cells treated with 1uM TriC, we observed a dose-dependent increase in intracellular BAX protein levels at 48 h, with sustained levels at 72 h of treatment. After 24 h of TriC treatment, MM.1S cells exhibited a robust transcriptional and metabolic response, consistent with the decreased viability, proliferation, and apoptosis observed at later time points. Indeed, many pro-apoptotic genes downstream of the ATF4-eIF2S1 pathway (42), such as *DDIT3* and *TRIB3,* showed an increasing trend of expression upon TriC treatment. Potentially related, two upstream activators of ATF4 the eIF2S1 pathway; PERK/EIF2AK3 and HRI/EIF2AK1, were enriched pathways.

Interestingly, the activation of ATF4 via PERK/EIF2AK3 or HRI/EIF2AK1 has been shown to induce apoptosis in AML (43) and MM (44,45). These data suggest that apoptosis may be initiated in a ATF4-eIF2S1-dependent manner and should be addressed in future studies.

TriC treatment for 24 h also significantly reduced MM.1S cell basal, maximal, and total ATP production rates due to a reduction in mitochondrially derived ATP. We observed a decrease in mitochondrial number and mitochondrial membrane potential after 24 h; therefore, it is unclear if the reduction in MM.1S cellular respiration and ATP production rates are definitively due to reduced mitochondria and/or compromised mitochondrial function. Our observation of suppressed superoxide levels at 24 h is consistent with an antioxidant transcriptional response, with *HMOX1* significantly increased after TriC treatment. As loss of mitochondrial function is associated with the production of superoxide, the subsequent increase in mitochondrial superoxide after 48 h of TriC treatment aligns with the proteomics data showing that proteins in the electron transport chain, such as NDUFV2, COX5A, COX6B1, and ATP5ME were significantly downregulated in TriC-treated MM.1S cells. However, only *COX5A* gene expression was significantly downregulated, suggesting that a post-translational mechanism may be involved.

Ferroptosis and lipid/atherosclerosis were among the most upregulated KEGG pathways in MM cells treated with 1µM TriC, as assessed by RNA-Seq. Additionally, the “Metabolism of Lipids” pathway was enriched in the upregulated genes, suggesting that pathways modulating or responding to lipid species within TriC-treated MM cells are activated. The majority of lipid metabolism-centered gene expression was related to anabolic processes, suggesting that MM.1S cells responded to lipid starvation, which is consistent with the predicted outcome of ACSL inhibition. Although we observed modest increases in apoptosis in response to 1 µM TriC treatment, it is possible that other mechanisms of cell death, including ferroptosis, which is characterized by the accumulation of lipid peroxides, play a role in the anti-MM action of TriC. It appears that both pro– and anti-ferroptotic genes have increased gene expression after 24 h of TriC treatment, suggesting that MM.1S cells are actively responding to TriC-dependent ferroptotic signals. Both ACSL3 and ACSL4 have been shown to regulate ferroptosis through independent mechanisms; thus, targeting both these proteins could have synergistic effects with ferroptosis agonists in multiple myeloma (8,46).

Combination therapies in MM are standard practice due to the development of drug resistance, and it is critical to identify new therapeutic combinations. TriC has been shown to synergize with other drugs, such as etoposide (19) and a combination of anti-metabolites and alkylating agents (47), in glioma and colorectal cancer. TriC has been shown to reduce lipid droplets (48), and lipid droplets have been identified as a mechanism of drug resistance in colorectal cancer (47). Together, these findings suggest that TriC treatment, in combination with other standard myeloma treatments, such as dexamethasone or proteasome inhibitors, may show promise. Additionally, MM cell Complex I and II activity is positively correlated with resistance to the BCL2 inhibitor, venetoclax (49). Hence, MM.1S cells treated with TriC may be sensitive to venetoclax treatment because we observed decreased mitochondrial ATP production and decreased expression of subunits of Complexes I and IV. In AML, AMPK-PERK-ATF4 activation also confers sensitivity to venetoclax treatment by repressing oxidative phosphorylation (43). Therefore, future studies should test whether a similar mechanism occurs in MM. Thus, targeting the ACSL family in combination with existing clinical treatments for MM is a promising prospect.

Our data supports the hypothesis that the ACSL family supports myeloma cell fitness. ACSLs represent a promising therapeutic target for MM, and further research is needed to elucidate the mechanisms by which this isozyme family supports MM cell survival, proliferation, and mitochondrial function. Future studies should investigate whether TriC and other metabolically targeted therapies could be combined with current myeloma treatments to achieve optimal anti-MM effects.

## Ethics Statement

This research did not meet the criteria for human subject research and did not use animal models.

## ACKNOWLEDGEMENTS

We thank all the multiple myeloma patients, their families, and healthcare providers for their Herculean efforts to keep fighting and the researchers who paved the road upon which this work was built. We thank Drs. Thomas Gridley, Benjamin King, Robert Koza, Lucy Liaw, and Clifford Rosen for reviewing the manuscript. We appreciate the help of Dr. Julie Dragon, Scott Tighe, Kristiaan Finstad, and all members of the Vermont Integrated Genomics Resource for their assistance with RNA-seq sample preparation. We thank Michele Karolak, Edward Jachimowicz and Dr. Yulica Santos-Ortega of the MHIR Molecular Phenotyping, Flow Cytometry and Proteomics and Lipidomics Cores, respectively. Thank you for the support CVM.

## AUTHOR CONTRIBUTIONS

**C.S. Murphy**: Conceptualization, resources, data curation, software, formal analysis, supervision, validation, investigation, visualization, methodology, writing-original draft, project administration, funding acquisition, writing-review, and editing. **V. DeMambro**: Resources, data curation, formal analysis, writing-original draft, writing–review, and editing. **S. Fadel**: Investigation. **H. Fairfield**: Investigation, project administration, funding acquisition, writing, reviewing, and editing. **C.A. Gartner**: LC-MS data acquisition, data curation, investigation, methodology, writing, review, and editing. **P. Rodriguez**: Data curation, investigation, writing, reviewing, and editing. **YW Qiang**: Investigation, writing-review, and editing **C. Vary**: curation, LC-MS data analysis, investigation, methodology, writing, reviewing, and editing. **M.R.Reagan**: Conceptualization, resources, data curation, formal analysis, supervision, funding acquisition, validation, investigation, visualization, methodology, writing – original draft, project administration, writing – review and editing

## Notes

Statement of Funding: We are thankful that this work was supported by the American Cancer Society (Research Grant RSG-19-037-01-LIB), NIH (R50CA265331, R37CA245330, R24 DK092759-01, P20GM121301, U54GM115516, and F31CA257695), and the Kane Foundation.

### Competing Interest Statement

The authors have declared no competing interest.

## REFERENCES

1. Siegel RL, Miller KD, Wagle NS, Jemal A. Cancer statistics, 2023. CA Cancer J Clin. 2023;73:17–48.

2. Hanahan D, Weinberg RA. Hallmarks of cancer: The next generation. Cell. 2011.

3. Roman-Trufero M, Auner HW, Edwards CM. Multiple myeloma metabolism – a treasure trove of therapeutic targets? Front Immunol. 2022;13:1–7.

4. Wang W, Zhao X, Wang H, Liang Y. Increased fatty acid synthase as a potential therapeutic target in multiple myeloma. J Zhejiang Univ Sci B [Internet]. 2008;9:441–7. Available from: http://www.springerlink.com/index/10.1631/jzus.B0740640

5. Tirado-Vélez JM, Joumady I, Sáez-Benito A, Cózar-Castellano I, Perdomo G. Inhibition of Fatty Acid Metabolism Reduces Human Myeloma Cells Proliferation. PLoS One. 2012;7.

6. Panaroni C, Fulzele K, Mori T, Onyewadume C, Raje NS. Multiple Myeloma Cells Induce Lipolysis in Adipocytes and Uptake Fatty Acids through Fatty Acid Transporter Proteins. Blood. 2020;136:41–2.

7. Farrell M, Fairfield H, Karam M, D’amico A, Murphy CS, Falank C, et al. Targeting the fatty acid binding proteins disrupts multiple myeloma cell cycle progression and MYC signaling. Elife. 2023;12:1–30.

8. Zhang J, Liu Y, Li Q, Zuo L, Zhang B, Zhao F, et al. ACSL4: a double-edged sword target in multiple myeloma, promotes cell proliferation and sensitizes cell to ferroptosis. Carcinogenesis [Internet]. 2023;44:242–51. Available from: 10.1093/carcin/bgad015

9. Quan J, Bode AM, Luo X. ACSL family: The regulatory mechanisms and therapeutic implications in cancer. Eur J Pharmacol [Internet]. Elsevier B.V.; 2021;909:174397. Available from: 10.1016/j.ejphar.2021.174397

10. Chen WC, Wang CY, Hung YH, Weng TY, Yen MC, Lai MD. Systematic analysis of gene expression alterations and clinical outcomes for long-chain acyl-coenzyme A synthetase family in cancer. PLoS One. 2016;11:1–23.

11. Sánchez-Martínez R, Cruz-Gil S, García-Álvarez MS, Reglero G, De Molina AR. Complementary ACSL isoforms contribute to a non-Warburg advantageous energetic status characterizing invasive colon cancer cells. Sci Rep. 2017;7:1–15.

12. Sánchez-Martínez R, Cruz-Gil S, de Cedrón MG, Álvarez-Fernández M, Vargas T, Molina S, et al. A link between lipid metabolism and epithelial-mesenchymal transition provides a target for colon cancer therapy. Oncotarget. 2015;6:38719–36.

13. Orlando UD, Castillo AF, Medrano MAR, Solano AR, Maloberti PM, Podesta EJ. Acyl-CoA synthetase-4 is implicated in drug resistance in breast cancer cell lines involving the regulation of energy-dependent transporter expression. Biochem Pharmacol [Internet]. Elsevier; 2019;159:52–63. Available from: 10.1016/j.bcp.2018.11.005

14. Wu X, Deng F, Li Y, Daniels G, Du X, Ren Q, et al. ACSL4 promotes prostate cancer growth, invasion and hormonal resistance. Oncotarget. 2015;6:44849–63.

15. Ma Y, Zha J, Yang XK, Li Q, Zhang Q, Yin A, et al. Long-chain fatty acyl-CoA synthetase 1 promotes prostate cancer progression by elevation of lipogenesis and fatty acid beta-oxidation. Oncogene [Internet]. Springer US; 2021;40:1806–20. Available from: 10.1038/s41388-021-01667-y

16. Wright HJ, Hou J, Xu B, Cortez M, Potma EO, Tromberg BJ, et al. CDCP1 drives triple-negative breast cancer metastasis through reduction of lipid-droplet abundance and stimulation of fatty acid oxidation. Proc Natl Acad Sci U S A. 2017;114:E6556–65.

17. Padanad MS, Konstantinidou G, Venkateswaran N, Melegari M, Rindhe S, Mitsche M, et al. Fatty Acid Oxidation Mediated by Acyl-CoA Synthetase Long Chain 3 Is Required for Mutant KRAS Lung Tumorigenesis. Cell Rep [Internet]. ElsevierCompany.; 2016;16:1614–28. Available from: 10.1016/j.celrep.2016.07.009

18. Igal RA, Wang P, Coleman RA. Triacsin C blocks de novo synthesis of glycerolipids and cholesterol esters but not recycling of fatty acid into phospholipid: evidence for functionally separate pools of acyl-CoA. Biochem J. Portland Press Ltd; 1997;324:529.

19. Mashima T, Sato S, Okabe S, Miyata S, Matsuura M, Sugimoto Y, et al. Acyl-CoA synthetase as a cancer survival factor: Its inhibition enhances the efficacy of etoposide. Cancer Sci. 2009;100:1556–62.

20. Ye W, Wang J, Huang J, He X, Ma Z, Li X, et al. ACSL5, a prognostic factor in acute myeloid leukemia, modulates the activity of Wnt/β-catenin signaling by palmitoylation modification. Front Med. 2023;

21. Chung SH, Lee HH, Kim YS, Song K, Kim TH. Induction of apoptosis in RL95-2 human endometrial cancer cells by combination treatment with docosahexaenoic acid and triacsin C. Arch Med Sci. Arch Med Sci; 2021;19:488–98.

22. Dempster JM, Boyle I, Vazquez F, Root DE, Boehm JS, Hahn WC, et al. Chronos: a cell population dynamics model of CRISPR experiments that improves inference of gene fitness effects. Genome Biol. Genome Biology; 2021;22:1–23.

23. Ghandi M, Huang FW, Jané-Valbuena J, Kryukov G V., Lo CC, McDonald ER, et al. Next-generation characterization of the Cancer Cell Line Encyclopedia. Nature [Internet]. Springer US; 2019;569:503–8. Available from: 10.1038/s41586-019-1186-3

24. Nchoutmboube JA, Viktorova EG, Scott AJ, Ford LA, Pei Z, Watkins PA, et al. Increased Long Chain acyl-Coa Synthetase Activity and Fatty Acid Import Is Linked to Membrane Synthesis for Development of Picornavirus Replication Organelles. PLoS Pathog. 2013;9.

25. Rappsilber J, Mann M, Ishihama Y. Protocol for micro-purification, enrichment, pre-fractionation and storage of peptides for proteomics using StageTips. Nat Protoc [Internet]. 2007;2:1896–906. Available from: 10.1038/nprot.2007.261

26. Fairfield H, Condruti R, Farrell M, Di Iorio R, Gartner CA, Vary C, et al. Development and characterization of three cell culture systems to investigate the relationship between primary bone marrow adipocytes and myeloma cells. Front Oncol. 2023;12:1–21.

27. Liberzon A, Birger C, Thorvaldsdóttir H, Ghandi M, Mesirov JP, Tamayo P. The Molecular Signatures Database Hallmark Gene Set Collection. Cell Syst. 2015;1:417–25.

28. Tomoda H, Kazuaki I, Satoshi O. Inhibition of acyl-CoA synthetase by triacsins. Biochim Biophys Acta (BBA)/Lipids Lipid Metab. 1987;921:595–8.

29. Love MI, Huber W, Anders S. Moderated estimation of fold change and dispersion for RNA-seq data with DESeq2. Genome Biol. 2014;15:1–21.

30. Croft D, Mundo AF, Haw R, Milacic M, Weiser J, Wu G, et al. The Reactome pathway knowledgebase. Nucleic Acids Res. 2014;42:472–7.

31. Kanehisa M, Furumichi M, Sato Y, Kawashima M, Ishiguro-Watanabe M. KEGG for taxonomy-based analysis of pathways and genomes. Nucleic Acids Res. Oxford University Press; 2023;51:D587–92.

32. Krämer A, Green J, Pollard J, Tugendreich S. Causal analysis approaches in ingenuity pathway analysis. Bioinformatics. 2014;30:523–30.

33. Yen MC, Kan JY, Hsieh CJ, Kuo PL, Hou MF, Hsu YL. Association of long-chain acyl-coenzyme A synthetase 5 expression in human breast cancer by estrogen receptor status and its clinical significance. Oncol Rep. 2017;37:3253–60.

34. Tomoda H, Igarashi K, Cyong JC, Omura S. Evidence for an essential role of long chain acyl-CoA synthetase in animal cell proliferation: Inhibition of long chain acyl-CoA synthetase by triacsins caused inhibition of Raji cell proliferation. J Biol Chem. 1991;266:4214–9.

35. Chung SH, Lee HH, Kim YS, Song K, Kim TH. Induction of apoptosis in RL95-2 human endometrial cancer cells by combination treatment with docosahexaenoic acid and triacsin C. Arch Med Sci. 2023;19:488–98.

36. Mashima T, Oh-hara T, Sato S, Mochizuki M, Sugimoto Y, Yamazaki K, et al. p53-defective tumors with a functional apoptosome-mediated pathway: A new therapeutic target. J Natl Cancer Inst. 2005;97:765–77.

37. Miller I, Min M, Yang C, Tian C, Gookin S, Carter D, et al. Ki67 is a Graded Rather than a Binary Marker of Proliferation versus Quiescence. Cell Rep [Internet]. ElsevierCompany.; 2018;24:1105–1112.e5. Available from: 10.1016/j.celrep.2018.06.110

38. Kim JH, Lewin TM, Coleman RA. Expression and characterization of recombinant rat acyl-CoA synthetases 1, 4, and 5: Selective inhibition by triacsin C and thiazolidinediones. J Biol Chem [Internet]. Â© 2001 ASBMB. Currently published by Elsevier Inc; originally published by American Society for Biochemistry and Molecular Biology.; 2001;276:24667–73. Available from: 10.1074/jbc.M010793200

39. Kaemmerer E, Peuscher A, Reinartz A, Liedtke C, Weiskirchen R, Kopitz J, et al. Human intestinal acyl-CoA synthetase 5 is sensitive to the inhibitor triacsin C. World J Gastroenterol. 2011;17:4883–9.

40. Al-Rashed F, Thomas R, Al-Roub A, Al-Mulla F, Ahmad R. LPS Induces GM-CSF Production by Breast Cancer MDA-MB-231 Cells via Long-Chain Acyl-CoA Synthetase 1. Molecules. Molecules; 2020;25.

41. Yu W, Cao D, Zhou H, Hu Y, Guo T. PGC-1α is responsible for survival of multiple myeloma cells under hyperglycemia and chemotherapy. Oncol Rep. 2015;33:2086–92.

42. Ohoka N, Yoshii S, Hattori T, Onozaki K, Hayashi H. TRB3, a novel ER stress-inducible gene, is induced via ATF4-CHOP pathway and is involved in cell death. EMBO J. 2005;24:1243–55.

43. Grenier A, Poulain L, Mondesir J, Jacquel A, Bosc C, Stuani L, et al. AMPK-PERK axis represses oxidative metabolism and enhances apoptotic priming of mitochondria in acute myeloid leukemia. Cell Rep [Internet]. The Authors; 2022;38:110197. Available from: 10.1016/j.celrep.2021.110197

44. Zhong Y, Zhang Y, Wang P, Gao H, Xu C, Li H. V8 induces apoptosis and the endoplasmic reticulum stress response in human multiple myeloma RPMI 8226 cells via the PERK-eIF2α-ATF4 signaling pathway. Oncol Lett. 2016;12:2702–9.

45. Burwick N, Zhang MY, de la Puente P, Azab AK, Hyun TS, Ruiz-Gutierrez M, et al. The eIF2-alpha kinase HRI is a novel therapeutic target in multiple myeloma. Leuk Res. England; 2017;55:23–32.

46. Klasson TD, LaGory EL, Zhao H, Huynh SK, Papandreou I, Moon EJ, et al. ACSL3 regulates lipid droplet biogenesis and ferroptosis sensitivity in clear cell renal cell carcinoma. Cancer Metab. 2022;10:1–17.

47. Cotte AK, Aires V, Fredon M, Limagne E, Derangère V, Thibaudin M, et al. Lysophosphatidylcholine acyltransferase 2-mediated lipid droplet production supports colorectal cancer chemoresistance. Nat Commun [Internet]. Springer US; 2018;9. Available from: 10.1038/s41467-017-02732-5

48. Fujimoto Y, Itabe H, Kinoshita T, Homma KJ, Onoduka J, Mori M, et al. Involvement of ACSL in local synthesis of neutral lipids in cytoplasmic lipid droplets in human hepatocyte HuH7. J Lipid Res [Internet]. Â© 2007 ASBMB. Currently published by Elsevier Inc; originally published by American Society for Biochemistry and Molecular Biology.; 2007;48:1280–92. Available from: 10.1194/jlr.M700050-JLR200

49. Bajpai R, Sharma A, Achreja A, Edgar CL, Wei C, Siddiqa AA, et al. Electron transport chain activity is a predictor and target for venetoclax sensitivity in multiple myeloma. Nat Commun [Internet]. Springer US; 2020;11. Available from: 10.1038/s41467-020-15051-z

50. Tang YUE, Zhou J, Hooi SC, Jiang YUEM, Lu GUOD. Fatty acid activation in carcinogenesis and cancer development: Essential roles of long – chain acyl – CoA synthetases (Review). 2018;1390–6.

