## Supplementary Materials for "Inhibition of Acyl-CoA Synthetase Long Chain Isozymes Decreases Multiple Myeloma Cell Proliferation and Causes Mitochondrial Dysfunction"

Conflict of Interest: All authors have no COIs.

Statement of Funding: We are thankful that this work was supported by the American Cancer Society (Research Grant RSG-19-037-01-LIB), NIH (R50CA265331, R37CA245330, R24 DK092759-01, P20GM121301, U54GM115516, and F31CA257695), and the Kane Foundation.

### Supplementary Data

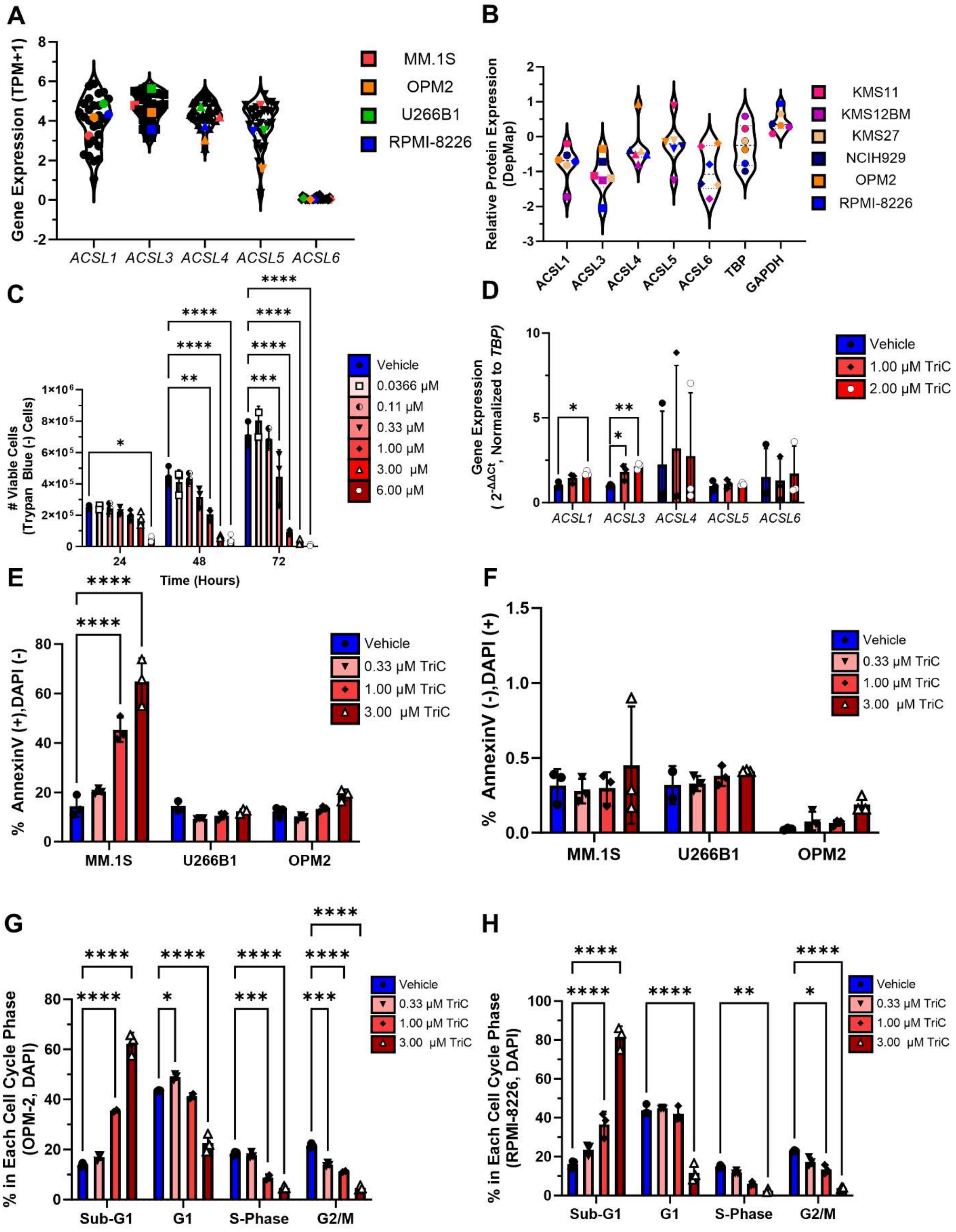

**Supplementary Figure 1: A,** Relative protein expression of the ACSL family and the house keeping proteins Tata-box binding protein (TBP) and glyceraldehyde-3-phosphate dehydrogenase (GAPDH) of 6 human myeloma cell lines from the Cancer Dependency Map/Cancer Cell Line Encyclopedia. **B,** ACSL family member gene expression (transcripts per million+1 (TPM+1) in 30 human myeloma cell lines from the Cancer Dependency Map/Cancer Cell Line Encyclopedia. Cell lines used in this study are highlighted. **C,** Expression of the ACSL family members in MM.1S cells treated with vehicle, 1, or 2 µM triacin C for 48 hours; assessed by qRT-PCR; n=3**. D-E,** Apoptosis (Annexin V-APC/DAPI) data for MM.1S, U266B1 and OPM2 treated with various doses of TriC for 48 hrs; n=3. Statistics: Panels **C**, **E-H**, Two-way ANOVA with Tukey's multiple comparisons test. **D,** One-way ANOVA with Dunnet's multiple comparisons test. Data are mean ± StDev, *p<0.05, **p<0.01, ***p<0.001 ****p<0.0001

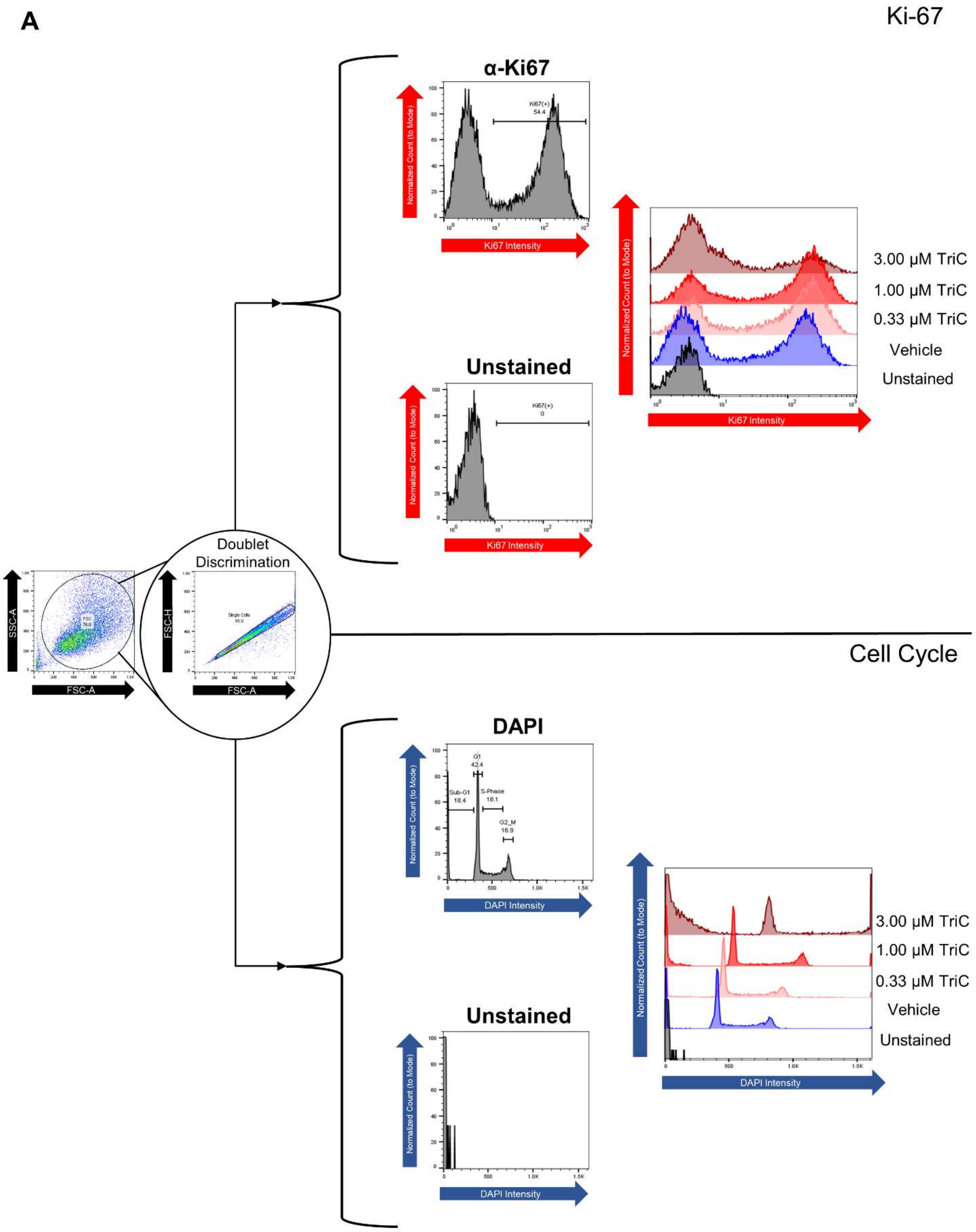
A
Ki-67
α-Ki67
Normalized Count (to Mode)
Ki67(+)
54.4
Ki67 Intensity
Unstained
Ki67(+)
0
3.00 µM TriC
1.00 µM TriC
0.33 µM TriC
Vehicle
Unstained
SSC-A
FSC-A
FSC
76.0
Doublet
Discrimination
FSC-H
Single Cells
95.0
Cell Cycle
DAPI
G1
42.4
Sub-G1
18.4
S-Phase
18.1
G2_M
16.9
DAPI Intensity

**Supplementary Figure 2: A,** Representative Ki-67 and cell cycle distribution flow cytometry plots depicting the gating strategies in MM.1S cells treated with various concentrations of TriC for 48 hours. An initial gate was made in the FSC-A vs. SSC-A and doublets were excluded comparing the FSC-A vs. FSC-H. In the same sample, both DAPI and Ki-67-AF647 was analyzed, positive populations were identified by comparing stained and unstained samples. Representative histograms of Ki-67 (top) and DAPI staining (bottom) depicting fluorescent intensities vs. normalized counts (to the mode) for various concentrations of TriC. A minimum of 10,000 events were collected.

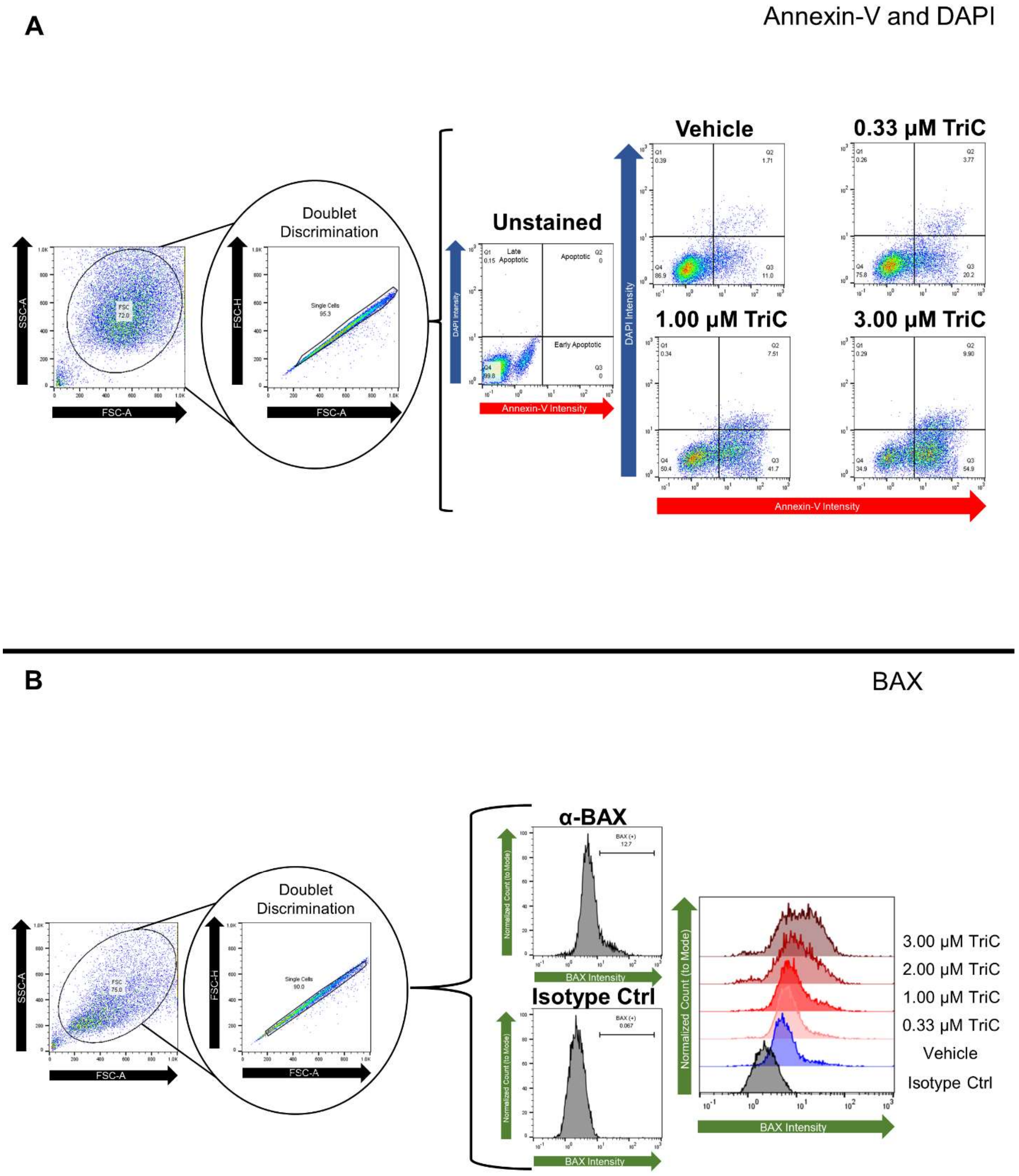


**Supplementary Figure 3: A,** Representative Apoptosis assay (Annexin V/DAPI) flow cytometry plots depicting the gating strategies in MM.1S cells treated with various concentrations of TriC for 48 hours. An initial gate was made in the FSC-A vs. SSC-A and doublets were excluded comparing the FSC-A vs. FSC-H. Within the single cell gate, positive populations were identified by comparing stained and unstained samples. Representative flow plots depicting fluorescent intensities of Annexin V vs. DAPI for various concentrations of TriC. A minimum of 10,000 events were collected. **B,** Representative BAX protein flow cytometry plots gated with a similar strategy as above, however positive signal was identified by comparing AF488 anti-BAX stained samples to AF488 Isotype control stained samples. A minimum of 10,000 events were collected.

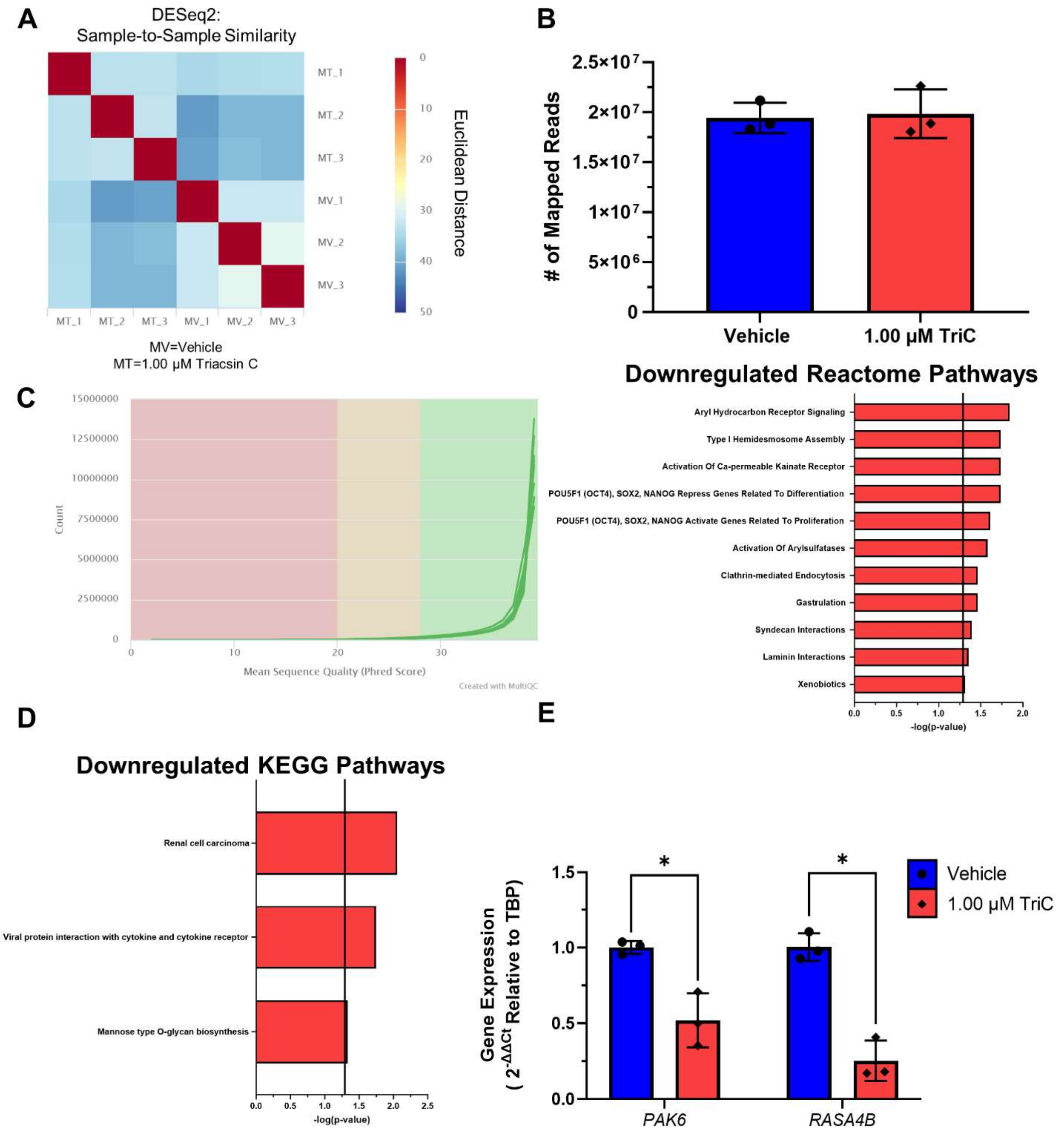


**Supplementary Figure 4: A,** RNA-Seq sample-to-sample similarity as displayed via Euclidean distances of the transcriptional profiles of MM.1S cells treated either with vehicle (DMSO, MV) or 1 µM triacsin C (TriC; MT) were calculated with DESeq2. Numbers associated with conditions designate different replicates. **B,** The number of reads mapped for MM.1S cells treated either with vehicle (DMSO) or 1 µM TriC after 24hrs. Reads were aligned to the *Homo sapiens* hg38 reference genome using STAR v2.7.10a and SAMtools v1.15.1, read counts were quantified using SALMON v1.5.250. **C,** The mean sequence quality (Phred Score) for each sample and their associated read counts are displayed for MM.1S cells treated either with vehicle (DMSO) or 1 µM triacsin for 24 hrs. Raw reads were subjected to quality checking and reporting (FastQC v0.11.9/ MultiQC v1.13[46]; and low quality sequence (Phred score <20) using Trim Galore v 0.6.7[47] were removed. **D-E,** Reactome and KEGG Pathways (respectively) associated with the significantly downregulated transcripts in 1 µM triacsin C treated MM.1S cells as assessed with Enrichr. **F,** Gene expression of significantly downregulated genes in the RNA-seq in MM.1S cells treated with vehicle or 1.00 µM triacsin C for 24hrs by qRT-PCR. Values are relative to *TBP*, n=3. **Statistics:** Unpaired Student's t-test or Welch's t-test. All data are mean ± StDev, *p<0.05, **p<0.01, ***p<0.001 ****p<0.0001

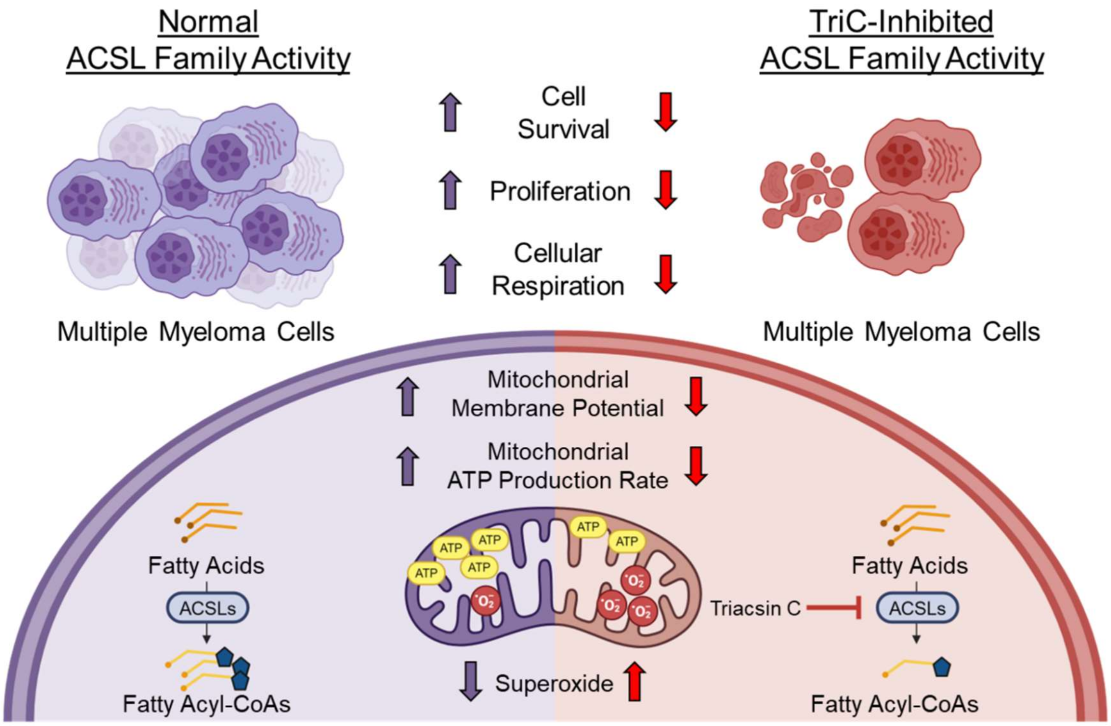


Triacsin C inhibition of the acyl-CoA synthetase long chain (ACSL) family decreases multiple myeloma cell survival, proliferation and reduces mitochondrial respiration and membrane potential.

Graphical Abstract made with Biorender.com

**Supplementary Table 1-qRT-PCR Forward Primers**

| Ensembl ID | Target Gene Name | Forward Primer (5'-3') | Tm (C) |
|---|---|---|---|
| ENSG00000160179 | ABCG1 | GTCTCGCTGATGAAAGGGCT | 60.11 |
| ENSG00000278540 | ACACA | ACAACGCAGGCATCAGAAGA | 57 |
| ENSG00000151726 | ACSL1 | GTGGAACTACAGGCAACCCC | 58.2 |
| ENSG00000123983 | ACSL3 | GGAACAATTTCCGAAGTGTGGG | 56.5 |
| ENSG00000068366 | ACSL4 | CCGCCCCTCCGCACAATAA | 60.6 |
| ENSG00000197142 | ACSL5 | TGCCAAAACCAAGTCAAAGCC | 56.8 |
| ENSG00000164398 | ACSL6 | AAATCGGCCAGAGTGGATCA | 56.4 |
| ENSG00000169020 | ATP5ME | GCCACGCGCTACAATTACCT | 61.09 |
| ENSG00000128965 | CHAC1 | GTGTGGAGGCCCGACTTC | 60.05 |
| ENSG00000178741 | COX5A | TTGATGCTCGCTGGGTAACA | 59.68 |
| ENSG00000126267 | COX6B1 | CGGGGTGCCTTTAGGATTCA | 59.75 |
| ENSG00000175197 | DDIT3 | GAGCTGGAAGCCTGGTATGA | 59.17 |
| ENSG00000168209 | DDIT4 | GGTTTGACCGCTCCACGAG | 61.03 |
| ENSG00000086232 | EIF2AK1 | CAACTCCGGGGTCCGCAA | 62.32 |
| ENSG00000172071 | EIF2AK3 | GCCAATTCAATGCCTGGGAC | 59.82 |
| ENSG00000128829 | EIF2AK4 | ATAACAAGCCCCCTCCCAAG | 59.37 |
| ENSG00000178607 | ERN1 | ACCCAGAGAAGCACGAAGAC | 59.68 |
| ENSG00000167468 | GPX4 | CAGTGAGGCAAGACCGAAGT | 59.97 |
| ENSG00000178127 | NDUFV2 | CCCGCCATGTTCTTCTCCG | 60.82 |
| ENSG00000188747 | NOXA1 | TGTGGATCGTGGGGACTGG | 61.29 |
| ENSG00000137843 | PAK6 | TCCAGCCCATGAAGACAGTG | 59.67 |
| ENSG00000158828 | PINK1 | CCTCCAGACGTGAGACAGTT | 59.04 |
| ENSG00000109819 | PPARGC1A | CCAAAGGATGCGCTCTCGTTCA | 63.42 |
| ENSG00000155846 | PPARGC1B | GGCGCTTTGAAGTGTTTGGT | 59.9 |
| ENSG00000087074 | PPP1R15A | CCCTAAAGGCCAGAAAGGTGC | 61.23 |
| ENSG00000170667 | RASA4B | ATCGTGGAGGGGAAGAACCT | 60.25 |
| ENSG00000099194 | SCD | GCTGTCAAAGAGAAGGGGAGT | 59.65 |
| ENSG00000168003 | SLC3A2 | GATGGGTTCCAGGTTCGGG | 60.08 |
| ENSG00000151012 | SLC7A11 | TGTGCTGACAAATGTGGCCT | 60.47 |
| ENSG00000112592 | TBP | GTGGGGAGCTGTGATGTGAA | 59.6 |
| ENSG00000108064 | TFAM | GCTCAGAACCCAGATGCAAAA | 59.11 |
| ENSG00000141510 | TP53 | TCAGATAGCGATGGTCTGGC | 59.32 |
| ENSG00000101255 | TRIB3 | TTCGCTGACCGTGAGAGGAAG | 62.08 |

**Supplementary Table 2-qRT-PCR Reverse Primers**

| Ensembl ID | Target Gene Name | Reverse Primer (5'-3') | Tm (C) | Product Length (bp) |
|---|---|---|---|---|
| ENSG00000160179 | ABCG1 | TGACTCAGGACGTAAAGCTGG | 59.73 | 110 |
| ENSG00000278540 | ACACA | GTTTCACCGCACACTGTTCC | 56.9 | 92 |
| ENSG00000151726 | ACSL1 | ATCATCTGGGCAAGGATTGAC | 55 | 113 |
| ENSG00000123983 | ACSL3 | CCCTGGGGTGTGGCTTATC | 58.2 | 127 |
| ENSG00000068366 | ACSL4 | ACAAGTGGACAGGCAGCAAAA | 57.7 | 129 |
| ENSG00000197142 | ACSL5 | TGTTGGTGTCAAGAGCCCAT | 56.8 | 75 |
| ENSG00000164398 | ACSL6 | TCATAGAGCGGGACCACCA | 58 | 72 |
| ENSG00000169020 | ATP5ME | TCTCTGGCAATCCGTTTCAGT | 59.65 | 101 |
| ENSG00000128965 | CHAC1 | ACACGGCCAGGCATCTTG | 60.36 | 119 |
| ENSG00000178741 | COX5A | ACAAGTGTGTTTATCCCTTTACGC | 59.79 | 79 |
| ENSG00000126267 | COX6B1 | GGGGCGGTCTTGTAGTTCTT | 59.68 | 72 |
| ENSG00000175197 | DDIT3 | GGTGAAGATTTTTGATTCTTCCTCT | 57.54 | 111 |
| ENSG00000168209 | DDIT4 | GGTAAGCCGTGTCTTCCTCC | 60.11 | 93 |
| ENSG00000086232 | EIF2AK1 | TTCTGCTGGAACATCAGATTCGTC | 61.15 | 127 |
| ENSG00000172071 | EIF2AK3 | TCCCGAGCCAATTCCCTATTG | 59.86 | 120 |
| ENSG00000128829 | EIF2AK4 | GCAGGATTTCACGTTGCTCC | 59.83 | 120 |
| ENSG00000178607 | ERN1 | GCTCCAGAAGAACGGGTGTT | 60.25 | 116 |
| ENSG00000167468 | GPX4 | TTACTCCCTGGCTCCTGCTT | 60.55 | 124 |
| ENSG00000178127 | NDUFV2 | ATTCCTTACATGTCTTCCCCAGT | 59.15 | 78 |
| ENSG00000188747 | NOXA1 | GGCTTGGTCAAATGCCCGC | 62.65 | 142 |
| ENSG00000137843 | PAK6 | AGGGTGTTGGAGCTGATGAC | 59.67 | 109 |
| ENSG00000158828 | PINK1 | CTCGGGCAGATGGTCTCTTG | 60.18 | 70 |
| ENSG00000109819 | PPARGC1A | CGGTGTCTGTAGTGGCTTGACT | 62.23 | 147 |
| ENSG00000155846 | PPARGC1B | CCGTACTTCTCGCCTCTCCT | 60.75 | 73 |
| ENSG00000087074 | PPP1R15A | TGCGATCCCGAGCAAGC | 60.18 | 120 |
| ENSG00000170667 | RASA4B | CACTGTGGCTGTCCTGATGA | 59.68 | 105 |
| ENSG00000099194 | SCD | AGCCAGGTTTGTAGTACCTCCT | 60.49 | 91 |
| ENSG00000168003 | SLC3A2 | CGCAATCAAGAGCCTGTCTTC | 59.6 | 108 |
| ENSG00000151012 | SLC7A11 | CGCTCAGAAAAGGTCACTGC | 59.49 | 87 |
| ENSG00000112592 | TBP | TGCTCTGACTTTAGCACCTGT | 59.31 | 183 |
| ENSG00000108064 | TFAM | GCCACTCCGCCCTATAAGC | 60.3 | 115 |
| ENSG00000141510 | TP53 | CTCATAGGGCACCACCACAC | 60.39 | 117 |
| ENSG00000101255 | TRIB3 | TTGTCCCACAGGGAATCATCTG | 60.03 | 89 |

**Supplementary Table 3- Average Chronos Scores of Modified Hallmark Fatty Acid Metabolism Genes in 21 Human Myeloma Cell Lines from the Cancer Dependency Map**

| Ensembl Gene ID | Gene Symbol | Gene Name | Avg Chronos Score | Std Dev |
|---|---|---|---|---|
| ENSG00000204370 | SDHD | succinate dehydrogenase complex subunit D | -1.435 | 0.244 |
| ENSG00000143252 | SDHC | succinate dehydrogenase complex subunit C | -1.420 | 0.266 |
| ENSG00000112972 | HMGCS1 | 3-hydroxy-3-methylglutaryl-CoA synthase 1 | -1.230 | 0.379 |
| ENSG00000164032 | H2AZ1 | H2A.Z variant histone 1 | -1.201 | 0.334 |
| ENSG00000164687 | FABP5 | fatty acid binding protein 5 | -1.118 | 0.308 |
| ENSG00000198856 | OSTC | oligosaccharyltransferase complex non-catalytic subunit | -1.034 | 0.239 |
| ENSG00000126088 | UROD | uroporphyrinogen decarboxylase | -0.928 | 0.318 |
| ENSG00000073578 | SDHA | succinate dehydrogenase complex flavoprotein subunit A | -0.896 | 0.280 |
| ENSG00000100412 | ACO2 | aconitase 2 | -0.854 | 0.338 |
| ENSG00000092010 | PSME1 | proteasome activator subunit 1 | -0.852 | 0.283 |
| ENSG00000119689 | DLST | dihydrolipoamide S-succinyltransferase | -0.799 | 0.172 |
| ENSG00000080819 | CPOX | coproporphyrinogen oxidase | -0.704 | 0.250 |
| ENSG00000025770 | NCAPH2 | non-SMC condensin II complex subunit H2 | -0.614 | 0.368 |
| ENSG00000091140 | DLD | dihydrolipoamide dehydrogenase | -0.564 | 0.250 |
| ENSG00000183955 | KMT5A | lysine methyltransferase 5A | -0.530 | 0.226 |
| ENSG00000072506 | HSD17B10 | hydroxysteroid 17-beta dehydrogenase 10 | -0.521 | 0.414 |
| ENSG00000099194 | SCD | stearoyl-CoA desaturase | -0.486 | 0.297 |
| ENSG00000188690 | UROS | uroporphyrinogen III synthase | -0.365 | 0.201 |
| ENSG00000158473 | CD1D | CD1d molecule | -0.337 | 0.196 |
| ENSG00000160124 | MIX23 | mitochondrial matrix import factor 23 | -0.328 | 0.170 |
| ENSG00000068366 | ACSL4 | acyl-CoA synthetase long chain family member 4 | -0.314 | 0.361 |
| ENSG00000091483 | FH | fumarate hydratase | -0.298 | 0.144 |
| ENSG00000278540 | ACACA | acetyl-CoA carboxylase alpha | -0.295 | 0.234 |
| ENSG00000146701 | MDH2 | malate dehydrogenase 2 | -0.269 | 0.231 |
| ENSG00000117592 | PRDX6 | peroxiredoxin 6 | -0.264 | 0.174 |
| ENSG00000102172 | SMS | spermine synthase | -0.251 | 0.212 |
| ENSG00000123983 | ACSL3 | acyl-CoA synthetase long chain family member 3 | -0.229 | 0.274 |
| ENSG00000124370 | MCEE | methylmalonyl-CoA epimerase | -0.218 | 0.115 |
| ENSG00000164024 | METAP1 | methionyl aminopeptidase 1 | -0.198 | 0.212 |

| | | | | |
|---|---|---|---|---|
| ENSG00000110090 | CPT1A | carnitine palmitoyltransferase 1A | -0.188 | 0.106 |
| ENSG00000128245 | YWHAH | tyrosine 3-monooxygenase/tryptophan 5-monooxygenase activation protein eta | -0.181 | 0.107 |
| ENSG00000122971 | ACADS | acyl-CoA dehydrogenase short chain | -0.169 | 0.188 |
| ENSG00000115255 | REEP6 | receptor accessory protein 6 | -0.160 | 0.130 |
| ENSG00000157184 | CPT2 | carnitine palmitoyltransferase 2 | -0.160 | 0.121 |
| ENSG00000205560 | CPT1B | carnitine palmitoyltransferase 1B | -0.160 | 0.111 |
| ENSG00000116882 | HAO2 | hydroxyacid oxidase 2 | -0.148 | 0.152 |
| ENSG00000131686 | CA6 | carbonic anhydrase 6 | -0.141 | 0.113 |
| ENSG00000163541 | SUCLG1 | succinate-CoA ligase GDP/ADP-forming subunit alpha | -0.136 | 0.168 |
| ENSG00000133835 | HSD17B4 | hydroxysteroid 17-beta dehydrogenase 4 | -0.130 | 0.152 |
| ENSG00000151726 | ACSL1 | acyl-CoA synthetase long chain family member 1 | -0.127 | 0.137 |
| ENSG00000083123 | BCKDHB | branched chain keto acid dehydrogenase E1 subunit beta | -0.122 | 0.138 |
| ENSG00000167315 | ACAA2 | acetyl-CoA acyltransferase 2 | -0.119 | 0.112 |
| ENSG00000198189 | HSD17B11 | hydroxysteroid 17-beta dehydrogenase 11 | -0.118 | 0.136 |
| ENSG00000169710 | FASN | fatty acid synthase | -0.118 | 0.199 |
| ENSG00000137106 | GRHPR | glyoxylate and hydroxypyruvate reductase | -0.117 | 0.099 |
| ENSG00000065833 | ME1 | malic enzyme 1 | -0.107 | 0.084 |
| ENSG00000161533 | ACOX1 | acyl-CoA oxidase 1 | -0.103 | 0.121 |
| ENSG00000109814 | UGDH | UDP-glucose 6-dehydrogenase | -0.103 | 0.185 |
| ENSG00000117305 | HMGCL | 3-hydroxy-3-methylglutaryl-CoA lyase | -0.103 | 0.100 |
| ENSG00000136143 | SUCLA2 | succinate-CoA ligase ADP-forming subunit beta | -0.101 | 0.134 |
| ENSG00000197375 | SLC22A5 | solute carrier family 22 member 5 | -0.100 | 0.087 |
| ENSG00000004961 | HCCS | holocytochrome c synthase | -0.100 | 0.123 |
| ENSG00000112033 | PPARD | peroxisome proliferator activated receptor delta | -0.099 | 0.134 |
| ENSG00000010932 | FMO1 | flavin containing dimethylaniline monoxygenase 1 | -0.095 | 0.114 |
| ENSG00000168291 | PDHB | pyruvate dehydrogenase E1 subunit beta | -0.083 | 0.205 |
| ENSG00000108515 | ENO3 | enolase 3 | -0.082 | 0.159 |
| ENSG00000095321 | CRAT | carnitine O-acetyltransferase | -0.082 | 0.102 |
| ENSG00000138413 | IDH1 | isocitrate dehydrogenase (NADP(+)) 1 | -0.081 | 0.169 |
| ENSG00000180902 | D2HGDH | D-2-hydroxyglutarate dehydrogenase | -0.080 | 0.129 |
| ENSG00000119471 | HSDL2 | hydroxysteroid dehydrogenase like 2 | -0.078 | 0.104 |
| ENSG00000138796 | HADH | hydroxyacyl-CoA dehydrogenase | -0.077 | 0.062 |
| ENSG00000104823 | ECH1 | enoyl-CoA hydratase 1 | -0.077 | 0.107 |
| ENSG00000080824 | HSP90AA1 | heat shock protein 90 alpha family class A member 1 | -0.071 | 0.113 |

| | | | | |
|---|---|---|---|---|
| ENSG00000148090 | AUH | AU RNA binding methylglutaconyl-CoA hydratase | -0.071 | 0.110 |
| ENSG00000240972 | MIF | macrophage migration inhibitory factor | -0.070 | 0.133 |
| ENSG00000106605 | BLVRA | biliverdin reductase A | -0.068 | 0.110 |
| ENSG00000104267 | CA2 | carbonic anhydrase 2 | -0.066 | 0.154 |
| ENSG00000101473 | ACOT8 | acyl-CoA thioesterase 8 | -0.066 | 0.113 |
| ENSG00000060971 | ACAA1 | acetyl-CoA acyltransferase 1 | -0.064 | 0.094 |
| ENSG00000111674 | ENO2 | enolase 2 | -0.060 | 0.131 |
| ENSG00000116791 | CRYZ | crystallin zeta | -0.056 | 0.090 |
| ENSG00000198130 | HIBCH | 3-hydroxyisobutyryl-CoA hydrolase | -0.053 | 0.088 |
| ENSG00000213316 | LTC4S | leukotriene C4 synthase | -0.051 | 0.202 |
| ENSG00000140465 | CYP1A1 | cytochrome P450 family 1 subfamily A member 1 | -0.051 | 0.068 |
| ENSG00000143819 | EPHX1 | epoxide hydrolase 1 | -0.050 | 0.105 |
| ENSG00000144724 | PTPRG | protein tyrosine phosphatase receptor type G | -0.049 | 0.168 |
| ENSG00000005187 | ACSM3 | acyl-CoA synthetase medium chain family member 3 | -0.047 | 0.144 |
| ENSG00000115361 | ACADL | acyl-CoA dehydrogenase long chain | -0.046 | 0.090 |
| ENSG00000139112 | GABARAPL1 | GABA type A receptor associated protein like 1 | -0.042 | 0.113 |
| ENSG00000241644 | INMT | indolethylamine N-methyltransferase | -0.040 | 0.122 |
| ENSG00000186951 | PPARA | peroxisome proliferator activated receptor alpha | -0.039 | 0.084 |
| ENSG00000104320 | NBN | nibrin | -0.036 | 0.109 |
| ENSG00000138029 | HADHB | hydroxyacyl-CoA dehydrogenase trifunctional multienzyme complex subunit beta | -0.035 | 0.103 |
| ENSG00000167434 | CA4 | carbonic anhydrase 4 | -0.033 | 0.093 |
| ENSG00000164398 | ACSL6 | acyl-CoA synthetase long chain family member 6 | -0.031 | 0.121 |
| ENSG00000172340 | SUCLG2 | succinate-CoA ligase GDP-forming subunit beta | -0.031 | 0.089 |
| ENSG00000176194 | CIDEA | cell death inducing DFFA like effector a | -0.030 | 0.140 |
| ENSG00000074416 | MGLL | monoglyceride lipase | -0.028 | 0.071 |
| ENSG00000042445 | RETSAT | retinol saturase | -0.027 | 0.080 |
| ENSG00000134333 | LDHA | lactate dehydrogenase A | -0.027 | 0.144 |
| ENSG00000167969 | ECI1 | enoyl-CoA delta isomerase 1 | -0.026 | 0.111 |
| ENSG00000198721 | ECI2 | enoyl-CoA delta isomerase 2 | -0.024 | 0.116 |
| ENSG00000170835 | CEL | carboxyl ester lipase | -0.020 | 0.099 |
| ENSG00000164434 | FABP7 | fatty acid binding protein 7 | -0.017 | 0.106 |
| ENSG00000131828 | PDHA1 | pyruvate dehydrogenase E1 subunit alpha 1 | -0.013 | 0.258 |
| ENSG00000115159 | GPD2 | glycerol-3-phosphate dehydrogenase 2 | -0.012 | 0.067 |

| | | | | |
|---|---|---|---|---|
| ENSG00000135821 | GLUL | glutamate-ammonia ligase | -0.012 | 0.131 |
| ENSG00000170323 | FABP4 | fatty acid binding protein 4 | -0.009 | 0.120 |
| ENSG00000078804 | TP53INP2 | tumor protein p53 inducible nuclear protein 2 | -0.006 | 0.117 |
| ENSG00000105679 | GAPDHS | glyceraldehyde-3-phosphate dehydrogenase, spermatogenic | -0.005 | 0.091 |
| ENSG00000012660 | ELOVL5 | ELOVL fatty acid elongase 5 | -0.004 | 0.069 |
| ENSG00000167588 | GPD1 | glycerol-3-phosphate dehydrogenase 1 | -0.001 | 0.084 |
| ENSG00000197142 | ACSL5 | acyl-CoA synthetase long chain family member 5 | 0.000 | 0.070 |
| ENSG00000104951 | IL4I1 | interleukin 4 induced 1 | 0.001 | 0.096 |
| ENSG00000109576 | AADAT | aminoadipate aminotransferase | 0.003 | 0.062 |
| ENSG00000104325 | DECR1 | 2,4-dienoyl-CoA reductase 1 | 0.005 | 0.093 |
| ENSG00000137274 | BPHL | biphenyl hydrolase like | 0.007 | 0.107 |
| ENSG00000147383 | NSDHL | NAD(P) dependent steroid dehydrogenase-like | 0.009 | 0.126 |
| ENSG00000123689 | G0S2 | G0/G1 switch 2 | 0.009 | 0.151 |
| ENSG00000115758 | ODC1 | ornithine decarboxylase 1 | 0.009 | 0.143 |
| ENSG00000197416 | FABP12 | fatty acid binding protein 12 | 0.011 | 0.147 |
| ENSG00000101365 | IDH3B | isocitrate dehydrogenase (NAD(+)) 3 non-catalytic subunit beta | 0.012 | 0.198 |
| ENSG00000089248 | ERP29 | endoplasmic reticulum protein 29 | 0.015 | 0.112 |
| ENSG00000113790 | EHHADH | enoyl-CoA hydratase and 3-hydroxyacyl CoA dehydrogenase | 0.017 | 0.091 |
| ENSG00000065057 | NTHL1 | nth like DNA glycosylase 1 | 0.018 | 0.114 |
| ENSG00000139547 | RDH16 | retinol dehydrogenase 16 | 0.018 | 0.090 |
| ENSG00000135218 | CD36 | CD36 molecule (CD36 blood group) | 0.020 | 0.094 |
| ENSG00000156587 | UBE2L6 | ubiquitin conjugating enzyme E2 L6 | 0.020 | 0.116 |
| ENSG00000121769 | FABP3 | fatty acid binding protein 3 | 0.020 | 0.049 |
| ENSG00000112299 | VNN1 | vanin 1 | 0.020 | 0.068 |
| ENSG00000127884 | ECHS1 | enoyl-CoA hydratase, short chain 1 | 0.021 | 0.119 |
| ENSG00000132196 | HSD17B7 | hydroxysteroid 17-beta dehydrogenase 7 | 0.023 | 0.123 |
| ENSG00000072778 | ACADVL | acyl-CoA dehydrogenase very long chain | 0.024 | 0.100 |
| ENSG00000154930 | ACSS1 | acyl-CoA synthetase short chain family member 1 | 0.025 | 0.098 |
| ENSG00000163586 | FABP1 | fatty acid binding protein 1 | 0.027 | 0.096 |
| ENSG00000151790 | TDO2 | tryptophan 2,3-dioxygenase | 0.029 | 0.073 |
| ENSG00000166228 | PCBD1 | pterin-4 alpha-carbinolamine dehydratase 1 | 0.030 | 0.083 |
| ENSG00000159231 | CBR3 | carbonyl reductase 3 | 0.030 | 0.178 |
| ENSG00000138696 | BMPR1B | bone morphogenetic protein receptor type 1B | 0.036 | 0.157 |
| ENSG00000159228 | CBR1 | carbonyl reductase 1 | 0.042 | 0.113 |

| | | | | |
|---|---|---|---|---|
| ENSG00000138698 | RAP1GDS1 | Rap1 GTPase-GDP dissociation stimulator 1 | 0.042 | 0.068 |
| ENSG00000170231 | FABP6 | fatty acid binding protein 6 | 0.043 | 0.065 |
| ENSG00000119673 | ACOT2 | acyl-CoA thioesterase 2 | 0.044 | 0.090 |
| ENSG00000169169 | CPT1C | carnitine palmitoyltransferase 1C | 0.046 | 0.109 |
| ENSG00000111897 | SERINC1 | serine incorporator 1 | 0.047 | 0.108 |
| ENSG00000150787 | PTS | 6-pyruvoyltetrahydropterin synthase | 0.053 | 0.091 |
| ENSG00000196344 | ADH7 | alcohol dehydrogenase 7 (class IV), mu or sigma polypeptide | 0.055 | 0.071 |
| ENSG00000100097 | LGALS1 | galectin 1 | 0.057 | 0.087 |
| ENSG00000189221 | MAOA | monoamine oxidase A | 0.060 | 0.084 |
| ENSG00000120694 | HSPH1 | heat shock protein family H (Hsp110) member 1 | 0.060 | 0.158 |
| ENSG00000132170 | PPARG | peroxisome proliferator activated receptor gamma | 0.070 | 0.104 |
| ENSG00000103150 | MLYCD | malonyl-CoA decarboxylase | 0.074 | 0.106 |
| ENSG00000100577 | GSTZ1 | glutathione S-transferase zeta 1 | 0.078 | 0.102 |
| ENSG00000197747 | S100A10 | S100 calcium binding protein A10 | 0.080 | 0.092 |
| ENSG00000067829 | IDH3G | isocitrate dehydrogenase (NAD(+)) 3 non-catalytic subunit gamma | 0.082 | 0.110 |
| ENSG00000105607 | GCDH | glutaryl-CoA dehydrogenase | 0.087 | 0.132 |
| ENSG00000145384 | FABP2 | fatty acid binding protein 2 | 0.088 | 0.071 |
| ENSG00000117054 | ACADM | acyl-CoA dehydrogenase medium chain | 0.088 | 0.111 |
| ENSG00000014641 | MDH1 | malate dehydrogenase 1 | 0.091 | 0.111 |
| ENSG00000162365 | CYP4A22 | cytochrome P450 family 4 subfamily A member 22 | 0.092 | 0.147 |
| ENSG00000067064 | IDI1 | isopentenyl-diphosphate delta isomerase 1 | 0.096 | 0.098 |
| ENSG00000120437 | ACAT2 | acetyl-CoA acetyltransferase 2 | 0.096 | 0.117 |
| ENSG00000171503 | ETFDH | electron transfer flavoprotein dehydrogenase | 0.104 | 0.118 |
| ENSG00000072042 | RDH11 | retinol dehydrogenase 11 | 0.105 | 0.112 |
| ENSG00000116133 | DHCR24 | 24-dehydrocholesterol reductase | 0.112 | 0.108 |
| ENSG00000187048 | CYP4A11 | cytochrome P450 family 4 subfamily A member 11 | 0.115 | 0.106 |
| ENSG00000164120 | HPGD | 15-hydroxyprostaglandin dehydrogenase | 0.122 | 0.124 |
| ENSG00000134240 | HMGCS2 | 3-hydroxy-3-methylglutaryl-CoA synthase 2 | 0.124 | 0.089 |
| ENSG00000136750 | GAD2 | glutamate decarboxylase 2 | 0.140 | 0.144 |

**Supplementary Table 4-Top 10 Significantly Upregulated Reactome Pathways in MM.1S Cells Treated with Triacsin C for 24 hours**

| Reactome Term | p-value | q-value | Significantly Associated Genes |
|---|---|---|---|
| Response Of EIF2AK1 (HRI) To Heme Deficiency R-HSA-9648895 | 1.16E-13 | 5.05E-11 | [PPP1R15A, CEBPB, DDIT3, ASNS, TRIB3, CHAC1, ATF3, ATF4] |
| Regulation Of Cholesterol Biosynthesis By SREBP (SREBF) R-HSA-1655829 | 1.66E-08 | 3.61E-06 | [SREBF1, HMGCS1, SCD, INSIG1, FASN, DHCR7, SEC24D, ACACA] |
| Cellular Responses To Stress R-HSA-2262752 | 1.45E-07 | 1.96E-05 | [PPP1R15A, KDM6B, JUN, CEBPB, CDKN1A, CBX4, EIF2AK3, ASNS, WIPI1, SLC7A11, FOS, HSPA13, VEGFA, ERN1, DDIT3, RPS6KA2, SESN2, HMOX1, TRIB3, CHAC1, ATF3, ATF4] |
| Cellular Responses To Stimuli R-HSA-8953897 | 2.02E-07 | 1.96E-05 | [PPP1R15A, KDM6B, JUN, CEBPB, CDKN1A, CBX4, EIF2AK3, ASNS, WIPI1, SLC7A11, FOS, HSPA13, VEGFA, ERN1, DDIT3, RPS6KA2, SESN2, HMOX1, TRIB3, CHAC1, ATF3, ATF4] |
| PERK Regulates Gene Expression R-HSA-381042 | 2.26E-07 | 1.96E-05 | [CEBPB, DDIT3, EIF2AK3, ASNS, ATF3, ATF4] |
| Unfolded Protein Response (UPR) R-HSA-381119 | 7.59E-07 | 5.49E-05 | [ERN1, CEBPB, DDIT3, EIF2AK3, ASNS, WIPI1, ATF3, ATF4] |
| Activation Of Gene Expression By SREBF (SREBP) R-HSA-2426168 | 1.22E-06 | 7.58E-05 | [SREBF1, HMGCS1, SCD, FASN, DHCR7, ACACA] |
| NR1H2 And NR1H3-mediated Signaling R-HSA-9024446 | 2.12E-06 | 0.0001 | [ABCA1, MYLIP, SREBF1, SCD, FASN, ABCG1] |
| ATF4 Activates Genes In Response To Endoplasmic Reticulum Stress R-HSA-380994 | 2.58E-06 | 0.0001 | [CEBPB, DDIT3, ASNS, ATF3, ATF4] |
| Metabolism Of Lipids R-HSA-556833 | 3.92E-05 | 0.0016 | [ABCA1, ARHGAP9, SREBF1, CHKB, HMGCS1, INSIG1, MID1IP1, ACSL3, SGPP2, ACACA, UGCG, SCD, FASN, |

| | | | |
|---|---|---|---|
| | | | LPCAT3, TRIB3, DHCR7, SEC24D, ABCD1] |

**Supplementary Table 5-Top 10 Significantly Upregulated KEGG Pathways in MM.1S Cells Treated with Triacsin C for 24 hours**

| KEGG Term | p-value | q-value | Significantly Associated Genes |
|---|---|---|---|
| Ferroptosis | 0.000001 | 0.000172 | [LPCAT3, HMOX1, SLC3A2, SLC7A11, ACSL3, SAT1] |
| Lipid and atherosclerosis | 0.000002 | 0.000172 | [ERN1, ABCA1, JUN, CCL3L1, DDIT3, EIF2AK3, CCL3, FOS, LDLR, ABCG1, ATF4] |
| Apoptosis | 0.000003 | 0.000178 | [ERN1, JUN, TUBA1A, DDIT3, EIF2AK3, PMAIP1, FOS, ATF4, BBC3] |
| Parathyroid hormone synthesis, secretion and action | 0.000029 | 0.001338 | [EGR1, CDKN1A, JUND, MAFB, FOS, ATF4, CREB5] |
| Fluid shear stress and atherosclerosis | 0.000163 | 0.005457 | [JUN, DUSP1, HMOX1, FOS, ACVR2A, ASS1, VEGFA] |
| MAPK signaling pathway | 0.000179 | 0.005457 | [JUN, JUND, DUSP10, DUSP1, DDIT3, RPS6KA2, FOS, DUSP16, ATF4, VEGFA] |
| Toll-like receptor signaling pathway | 0.000232 | 0.006055 | [JUN, CCL3L1, CCL4L2, CCL4, CCL3, FOS] |
| Non-alcoholic fatty liver disease | 0.000318 | 0.00727 | [ERN1, SREBF1, JUN, DDIT3, EIF2AK3, FOS, ATF4] |
| Fatty acid biosynthesis | 0.000418 | 0.008501 | [FASN, ACSL3, ACACA] |
| AMPK signaling pathway | 0.000500 | 0.008702 | [SREBF1, SCD, FASN, ACACA, PCK2, CREB5] |

**Supplementary Table 6-Top 10 Significantly Downregulated Reactome Pathways in MM.1S Cells Treated with Triacsin C for 24 hours**

| Reactome Term | p-value | q-value | Significantly Associated Genes |
|---|---|---|---|
| Aryl Hydrocarbon Receptor Signaling R-HSA-8937144 | 0.0142 | 0.3086 | [ARNT2] |
| POU5F1 (OCT4), SOX2, NANOG Repress Genes Related To Differentiation R-HSA-2892245 | 0.0183 | 0.3086 | [SOX2] |
| Activation Of Ca-permeable Kainate Receptor R-HSA-451308 | 0.0183 | 0.3086 | [GRIK4] |
| Type I Hemidesmosome Assembly R-HSA-446107 | 0.0183 | 0.3086 | [ITGB4] |
| POU5F1 (OCT4), SOX2, NANOG Activate Genes Related To Proliferation R-HSA-2892247 | 0.0243 | 0.3086 | [SOX2] |
| Activation Of Arylsulfatases R-HSA-1663150 | 0.0263 | 0.3086 | [ARSI] |
| Clathrin-mediated Endocytosis R-HSA-8856828 | 0.0342 | 0.3086 | [AMPH, FNBP1L] |
| Gastrulation R-HSA-9758941 | 0.0342 | 0.3086 | [SOX2] |
| Syndecan Interactions R-HSA-3000170 | 0.0402 | 0.3086 | [ITGB4] |
| Laminin Interactions R-HSA-3000157 | 0.0441 | 0.3086 | [ITGB4] |

**Supplementary Table 7- Significantly Downregulated KEGG Pathways in MM.1S Cells Treated with Triacsin C for 24 hours**

| KEGG Term | p-value | q-value | Significantly Associated Genes |
|---|---|---|---|
| Renal cell carcinoma | 0.0088 | 0.35041 | [ARNT2, PAK6] |
| Viral protein interaction with cytokine and cytokine receptor | 0.0178 | 0.35041 | [IL22RA1, CX3CR1] |
| Mannose type O-glycan biosynthesis | 0.0461 | 0.35041 | [B3GAT2] |

**Supplementary Table 8- Significantly Changed Proteins Shared Among Overrepresented Pathways between MM.1S Cells Treated with 1 or 2 µM Triacsin C for 48 hours**

| UniProt ID | Gene Symbol | 1 µM TriC p-value | 1 µM TriC log2(FC) | 2 µM TriC p-value | 2 µM TriC log2(FC) |
|---|---|---|---|---|---|
| P14854 | COX6B1 | 0.00036 | -2.4301 | 0.00196 | -3.1648 |
| P00441 | SOD1 | 0.00311 | -1.7038 | 0.0058 | -2.3216 |
| P30046 | DDT | 0.0025 | -1.649 | 0.00148 | -2.0939 |
| Q13404 | UBE2V1 | 0.00441 | -1.3851 | 0.00383 | -1.8907 |
| P14174 | MIF | 0.00439 | -1.3405 | 0.0057 | -2.0357 |
| P60174 | TPI1 | 0.0077 | -1.1878 | 4.3E-05 | -1.8122 |
| P56385 | ATP5ME | 0.03366 | -1.1303 | 0.00585 | -2.2481 |
| P25774 | CTSS | 0.02113 | -1.1126 | 8.6E-05 | -1.8819 |
| Q06323 | PSME1 | 0.01176 | -1.1088 | 0.00083 | -1.5907 |
| P20674 | COX5A | 0.01349 | -0.855 | 0.00256 | -0.8827 |
| P62136 | PPP1CA | 0.00324 | -0.7742 | 0.01815 | -0.8254 |
| Q9BWD1 | ACAT2 | 0.04915 | -0.7617 | 0.00058 | -0.9762 |
| Q9Y3D6 | FIS1 | 0.00941 | -0.7583 | 0.01764 | -0.7654 |
| P19404 | NDUFV2 | 0.00352 | -0.7359 | 0.01528 | -0.9045 |
| P63279 | UBE2I | 0.02546 | -0.715 | 0.01008 | -1.0345 |
| P14406 | COX7A2 | 0.0032 | -0.7146 | 0.02682 | -0.5887 |
| P30044 | PRDX5 | 0.0071 | -0.6596 | 0.00767 | -0.577 |
| P62979 | RPS27A | 0.03995 | -0.6181 | 0.01422 | -0.8281 |
| P32119 | PRDX2 | 0.02419 | -0.5751 | 0.00307 | -0.7292 |
| Q99497 | PARK7 | 0.03034 | -0.5744 | 0.00112 | -0.8881 |
| Q06830 | PRDX1 | 0.01014 | -0.5517 | 0.00579 | -0.6419 |
| P30041 | PRDX6 | 0.02302 | -0.4917 | 0.0104 | -0.5503 |
| P40926 | MDH2 | 0.00938 | -0.4814 | 0.00099 | -0.6454 |
| Q99459 | CDC5L | 0.04299 | -0.4597 | 0.00621 | -1.0905 |
| P67775 | PPP2CA | 0.0143 | -0.4411 | 0.01451 | -0.4875 |
| Q8NCW5 | NAXE | 0.03351 | -0.4206 | 0.04732 | -0.444 |
| Q12906 | ILF3 | 0.03873 | -0.3223 | 0.005 | -0.6077 |
| P05387 | RPLP2 | 0.03781 | -0.3036 | 0.01911 | -0.5576 |
| P27797 | CALR | 0.01821 | -0.2959 | 0.01418 | -0.3125 |
| P04406 | GAPDH | 0.04388 | -0.2738 | 0.00747 | -0.377 |
| Q8WXX5 | DNAJC9 | 0.00801 | -0.2674 | 0.0031 | -0.8543 |
| P17844 | DDX5 | 0.01053 | -0.233 | 0.01265 | -0.413 |
| P00505 | GOT2 | 0.03319 | -0.2079 | 0.0133 | -0.356 |
| P68371 | TUBB4B | 0.04662 | -0.1506 | 0.02135 | -0.1749 |
| P18124 | RPL7 | 0.04337 | 0.1003 | 0.03231 | 0.12958 |
| P46459 | NSF | 0.03706 | 0.16247 | 0.00657 | 0.27477 |
| P42765 | ACAA2 | 0.02503 | 0.19968 | 0.00956 | 0.2899 |

| | | | | | |
|---|---|---|---|---|---|
| P49770 | EIF2B2 | 0.00981 | 0.43057 | 0.00854 | 0.57185 |
| P50416 | CPT1A | 0.02552 | 1.02346 | 0.00218 | 1.05261 |
